# Quorum Sensing Modulates Bacterial Virulence and Colonization Dynamics During an Enteric Infection

**DOI:** 10.1101/2023.03.14.532656

**Authors:** Jorge Peña-Díaz, Sarah E. Woodward, Anna Creus-Cuadros, Antonio Serapio-Palacios, Wanyin Deng, B. Brett Finlay

**Affiliations:** Department of Microbiology and Immunology, University of British Columbia, Vancouver, BC, V6T 1Z3, Canada; Michael Smith Laboratories, University of British Columbia, Vancouver, BC, V6T 1Z4, Canada; Department of Biochemistry and Molecular Biology, University of British Columbia, Vancouver, BC, V6T 1Z3, Canada

**Keywords:** Quorum Sensing, bacterial communication, AHLs, autoinducers, chemical signaling, RNA-seq, bacterial attachment, virulence, type III secretion system, pathogenesis.

## Abstract

Quorum Sensing (QS) is a form of cell-to-cell communication that enables bacteria to modify behaviour according to their population density. While QS has been proposed as a potential intervention against pathogen infection, QS-mediated communication within the mammalian digestive tract remains understudied. Using an LC-MS/MS approach, we discovered that *Citrobacter rodentium*, a natural murine pathogen used to model human infection by pathogenic *Escherichia coli*, utilizes the CroIR system to produce three QS-molecules. We then profiled their accumulation both *in vitro* and across different gastrointestinal sites over the course of infection. Importantly, we found that in the absence of QS capabilities the virulence of *C. rodentium* is enhanced. This highlights the role of QS as an effective mechanism to regulate virulence according to the pathogen’s spatio-temporal context to optimize colonization and transmission success. These results also demonstrate that inhibiting QS may not always be an effective strategy for the control of virulence.

## Introduction

Bacteria are often found as members of complex heterogenous communities. To survive in diverse and competitive environments, such as within the gastrointestinal tract, microbes need to sense and adapt to a variety of environmental barriers such as host physiology, nutrient availability, osmotic stress, and the presence of other bacteria^1–3^. The use of chemical signals to perform cell-to-cell communication, a process known as quorum sensing (QS), enables bacteria to sense and respond to population density through the production of small diffusible molecules known as autoinducers (AIs)^4–6^. This process allows bacteria to coordinate behaviors at the population level which enables them to regulate the expression of metabolically expensive activities such as bioluminescence^7, 8^, expression of virulence genes^9, 10^ and biofilm formation^11–13^. This way, QS facilitates dynamic alterations of the bacterial transcriptome according to the surrounding environment and promotes rapid adaptation to different environmental niches to increase population fitness.

Many different mechanisms for QS have evolved among both bacterial pathogens and commensals, however the most common class of AIs in Gram-negative bacteria belongs to the family of N-acyl-homoserine lactones (AHLs). AHLs share a common homoserine-lactone ring, but differ in their structure either by the length of their acyl-chain, or by the presence or absence of a substitution (such as a hydroxy- or an oxo-group) at the position C3^14, 15^. These characteristics can significantly affect stability, signaling dynamics, and specificity to their cognate receptors, explaining why they are generally regarded as intra-species communication signals^16^. AHL-mediated QS systems are often composed of a LuxI-family AHL synthase, and a partner LuxR-family AHL response regulator. When a LuxR-type receptor comes in contact with its cognate AHL, they often form LuxR-AHL complexes. This interaction stabilizes the protein complex and facilitates its binding to specific DNA sequences known as *“lux* boxes”, where LuxR-type receptors can act as either transcriptional activators or repressors^15^. QS signaling networks can thereby translate information about the AI levels present in the environment into changes in bacterial gene expression.

Although QS has been described in a wealth of environments, QS-mediated communication within the mammalian digestive tract remains largely understudied. This is surprising given that QS is predicted to play a major role in environmental niches where complex communities of microorganisms interact. Several factors, including low accumulation of AIs and high susceptibility for degradation, make studying cell-to-cell communication in the gut environment exceptionally challenging^17^. Nevertheless, research focusing on QS within the gastrointestinal tract is rapidly becoming an area of great interest, where QS has already been shown to play an important role within the host context regulating processes such as maintenance of microbiota eubiosis^18^, immune regulation^19, 20^, or impeding pathogenicity of harmful bacteria^21^. Better understanding chemical communication within the gastrointestinal tract has great potential in developing innovative therapeutic interventions to selectively manipulate microbial behaviors and promote human health.

To better understand AHL-mediated QS signaling by gastrointestinal pathogens, we investigated its importance during natural infection by *Citrobacter rodentium*, a murine pathogen used to model human infections by enteropathogenic and enterohemorrhagic *Escherichia coli* (EPEC and EHEC)^22, 23^. During infection, *C. rodentium*, EPEC, and EHEC induce the formation of attaching and effacing (A/E) lesions, which are characterized by intimate bacterial attachment to the host epithelia, formation of actin rich-pedestals, and the destruction of the brush border microvilli ^24, 25^. This process is mediated by the type III secretion system (T3SS), a molecular syringe-like apparatus encoded by five gene operons within the Locus of Enterocyte Effacement (LEE1-5)^26–28^. A/E pathogens utilize the T3SS to directly inject effector proteins into the host cell to subvert various host processes. While the virulence machinery of *C. rodentium* has been extensively studied, little is known about the role that its LuxIR homolog system (CroIR) plays during host infection or the specific signaling dynamics that AHLs have in coordinating pathogen behavior.

In this study, we developed an LC-MS/MS method to characterize the production of AHLs by *C. rodentium* both *in vitro* and across different gastrointestinal sites over the course of infection. Our results revealed that disrupting the natural ability of *C. rodentium* to respond to its population density led to a significant increase in pathogenicity both *in vitro* and *in vivo*. We propose that the spatio-temporal composition of AHLs in the environment enables *C. rodentium* to optimize the timing for epithelial attachment in a T3SS-dependent manner, thereby enhancing pathogen fitness during infection and transmission. These findings also demonstrate that the interruption of QS signaling as a therapeutic strategy may not always be effective and should not be considered a generalized strategy for the control of virulence.

## Materials and methods

### Bacterial strains and culturing conditions

Bacterial strains and plasmids used in this study are listed in the Supplemental Tables S1-S2. Unless otherwise stated, bacterial strains were routinely grown aerobically in Lysogeny Broth (LB) at 37 °C with agitation at 220 rpm. When necessary, chloramphenicol was supplemented at 30 µg/mL.

### Construction of deletion and complemented strains

Chromosomal in-frame deletions of *croI* and *croR* were created via allelic exchange. In short, target deletion constructs were generated by using primers found in Supplemental Table S3. Purified DNA fragments were ligated into the suicide plasmid pRE112 by using Gibson assembly^29^. Biparental mating was performed using MFD*pir*^30^ as a donor strain to transfer the plasmid into *C. rodentium.* The loss of the pRE112 plasmid was induced via sucrose counter-selection^31^. In-frame deletion was confirmed via PCR and sequencing.

Chromosomally complemented strains were generated by amplifying the gene of interest along with its native promoter using primers listed in Supplemental Table S4. Constructs were then cloned into the pUC18R6KT-mini-Tn7T plasmid system^32^. Conjugation to the target *Citrobacter* strain was achieved via triparental mating using MDF*pir* as the donor and pTNS2^32^ as the helper strain. Correct chromosomal insertion was confirmed via PCR and sequencing by using the primers PTn7R and PglmS-down (Supplemental Table S4). All deletion and complemented strains had similar growth rates to wild-type (WT) (Figure S1).

### Materials and reagents

N-butyryl-L-homoserine lactone (C4-HSL), N-hexanoyl-L-homoserine lactone (C6-HSL) and N-(β-ketocaproyl)-L-homoserine lactone (3-oxo-C6-HSL) and N-tetradecanoyl-L-homoserine lactone (C14-HSL) were purchased from Cayman Chemical Company (Ann Arbor, MI, USA). 3-hydroxy-hexanoyl-L-homoserine lactone (3-hydroxy-C6-HSL) was obtained from Chemodex (St. Gallen, Switzerland). Analytical standards were resuspended in dimethyl sulfoxide (DMSO) and stored at −70 °C until analysis. HPLC-grade acetonitrile was obtained from Fisher Chemical; HPLC-grade ethyl acetate, formic acid (FA), and DMSO were purchased from Sigma-Aldrich (St. Louis, MO, USA).

### Mouse infections

Animal experiments were performed in accordance with the Canadian Council on Animal care and the University of British Columbia (UBC) Animal Care Committee guidelines (Animal Care Protocol A20-0187). Female C57BL/6J mice were purchased from Jackson Laboratory (Bar Harbor, ME) and maintained on a 12-hour light-dark cycle in a specific pathogen-free facility at UBC. All mice were allowed to acclimatize to the facility for a period of one week upon arrival.

*C. rodentium* infection was performed as follows: mice were gavaged orally with 10^8^ CFU (in 100 µL volume) of either the WT, or Δ*croI* strain of *C. rodentium* from overnight cultures, as confirmed by retrospective plating. Mice were monitored daily post-gavage for weight loss and clinical symptoms, before euthanasia at experimental endpoint on day 4 or day 7 post-infection by isoflurane anesthesia followed by carbon dioxide inhalation. Mice were 7 weeks old at the time of infection. Mice that remained uninfected on the day of the experimental endpoint were removed from all analyses.

To assess morbidity and mortality, female C3H/HeJ mice were purchased from Jackson Laboratory. At 7-weeks-old mice were gavaged orally with 10^8^ CFU of either the wild-type, mutant or complemented strains of *C. rodentium* (100 µL volume) from overnight cultures. Mice were monitored twice daily for weight loss and clinical symptoms, and were euthanized upon reaching humane endpoint.

### *In vivo* sample collection

Fecal samples were collected every 2 days post-infection to enumerate *C. rodentium* burden. At experimental endpoint, large intestinal (cecum and colon) and systemic (spleen) organs were collected. Cecum and colon samples were separated into lumenal and tissue-associated subpopulations by opening the tissues longitudinally to gently collect lumenal content, before washing the remaining tissue twice in phosphate buffered saline (PBS) for collection of mucosal associated-bacteria. Spleens were collected whole.

CFU enumeration of bacterial burden was performed by collecting samples in 1 mL PBS followed by homogenization in a FastPrep-24 (MP Biomedicals) at 5.5 m/s for 2 minutes. Sample homogenate was then diluted for plating on MacConkey agar (Difco). Bacterial plates were incubated at 37 °C for 18-20 hours before counting.

Intestinal samples for AHL quantification were collected from cecal and colonic contents as described above. Samples were weighed for normalization and stored at −70 °C until further processing during AHL extraction (see ‘AHL extraction and sample preparation’).

### Supernatants for *in vitro* detection of AHLs

Overnight cultures of *C. rodentium* were inoculated (1:40) in 5 mL of the different media of interest: LB broth and YCFAG (Yeast extract, Casitone and Fatty Acid Glucose^33^) were incubated with agitation at 37 °C; DMEM (Dulbecco’s Modified Eagle’s Medium; Hyclone), and DMEM+++ (DMEM supplemented with 10 % heat-inactivated fetal bovine serum, 1 % Non-Essential Amino Acids, and 1 % L-glutamine) were incubated statically at 37 °C in 5 % CO2. Cell-free bacterial-derived supernatants were obtained at 6-, 8-, 12- and 24-hours post-inoculation by centrifuging and filter sterilizing extracts through a 0.22 µm filter. Extracts were stored at −70 °C until processing.

### AHL extraction and sample preparation

AHLs were extracted by utilizing a method based on Shaw^34^ and Zhu^35^ with modifications. In short, for *in vitro* experiments, cell-free derived supernatants were extracted three times using 3 volumes of ethyl acetate. The organic fraction was collected and evaporated until dry using a vacuum centrifuge (Vacufuge, Eppendorf). Samples were resuspended in 200 µL of 70 % acetonitrile acidified with 0.1 % formic acid (FA), filtered through a 0.22 µm porosity filter, and stored at −70 °C until analysis. For murine intestine-derived samples, extractions were performed three times by first homogenizing cecal and colonic contents in 50 % ethyl acetate using a Mixer Mill MM 400 (Retsch), followed by constant agitation for at least 10 minutes at 4 °C using a rocker. Mixtures were centrifuged at 8000 ×*g* for 10 minutes, and the supernatants were collected into a separate tube. Samples were re-extracted three times in 100 % ethyl acetate, collecting only the organic fraction after centrifugation. Extracts were completely dried using a vacuum centrifuge, reconstituted in 50 µL of 70 % acetonitrile (0.1 % FA), filtered through a 0.22 µm porosity filter, and stored at −70 °C until analysis.

### LC-MS/MS analysis

Extracted samples were injected into an Agilent 6460 Triple Quadrupole (QQQ) Mass Spectrometer (Agilent) equipped with an electrospray ionization source operated in the positive ion mode. A reversed-phase Nucleosil C18 5 µm, 250 × 4.6 mm column was used for chromatographic separation of AHLs. MS was conducted with an electrospray ionization voltage of 3500 V, while using 50 psi nebulizer gas at a temperature of 350 °C. The sample injection volume was 5 µL. LC separation was performed while using mobile phase A (0.1 % FA in 3 % acetonitrile 97 % water) and mobile phase B (0.1 % FA in 90 % acetonitrile 10 % water), at a flow rate of 450 µL/min. The separation gradient was as follows: 40 % B for 2 min, 40 to 60 % B in 4 min, 60 to 70 % B in 2 min, 70 to 80 % in 3 min, 80 % for 1 min, 80 to 40 % in 1 min, and 40 % for 2 min. For the identification and quantification of analytes, multiple-reaction-monitoring (MRM) was utilized (Supplemental Table S6). Data were analyzed using MassHunter Qualitative Analysis B.06.00 software (Agilent Technologies). Calibration curves were constructed with a minimum of 6-points using analytical standards of C4-HSL, C6-HSL, 3-hydroxy-C6-HSL and 3-oxo-C6-HSL with concentrations ranging from 0.1 to 250 ppb for murine intestine-derived samples, and from 10 to 5000 ppb for *in vitro* experiments. Regression coefficients (*R*^2^) of all AHL-calibration curves were greater than 0.99.

### RNA isolation and sequencing

Wild-type and Δ*croI* strains were subcultured in 6-well plates with 3 mL of DMEM and grown statically for 6 hours at 37 °C in 5 % CO2. Bacterial cells were collected and treated with RNAprotect Bacteria Reagent (Qiagen) and stored at −70 °C until extraction. RNA was isolated using the Qiagen RNeasy Mini Kit, and genomic DNA was digested on column by using the RNase-Free DNase Set (Qiagen). RNA quality was assessed using a 2100 Bioanalyzer (Agilent Technologies, USA); all samples had RNA Integrity scores higher than 6 which were considered as high quality for *C. rodentium*^36^. Library preparation, rRNA depletion, and sequencing were performed by the Canada’s Michael Smith Genome Sciences Centre. In short, rRNA was depleted by using the bacterial NEBNext rRNA Depletion Kit (New England BioLabs, E7850). Sequencing libraries were created by end repair and phosphorylation, followed by 3’ A-addition and adapter ligation using a custom reagent formulation (New England BioLabs, E6000B-10). Libraries were pooled in equal molar amounts and were sequenced using an Illumina HiSeqX platform (Ilumina) using paired-end (PE) reads.

### RNA-seq data analysis workflow and pathway enrichment analysis

Reads were aligned to *C. rodentium* ICC168 reference genome (NC_013716, NC_013717, NC_013718 and NC_013719) by using STAR 2.7.10a^37^ (v2.7.10a). Quality of the alignment was assessed by using FastQC^38^ (v0.11.9), MultiQC^39^ (v1.12), and RSeQC^40^ (v4.0.0). At least 9 million PE reads were obtained from each library. General alignment and quality statistics are available in Supplementary Data 1. Gene counts were imported into R (v4.1.0) for further processing (Supplementary Data 2). Only reads that were present in at least 3 samples with a count of 10 or higher were included for analysis. Differentially expressed genes (DEGs) were identified via DESeq2^41^ using Independent Hypothesis Weighting (IHW) for *p*-value adjustment^42^ and ‘apeglm’ for log fold-change shrinkage^43^. Genes were considered differentially regulated when the comparison between Δ*croI* and WT passed an adjusted *p*-value of < 0.05 and had a fold change equal or greater than ± 1.25 (Supplementary Data 3).

Pathway enrichment analysis was performed by using Gene Set Enrichment Analysis (GSEA, v4.2.3)^44, 45^ using a pre-ranked list of all genes within the dataset (after pre-filtering) arranged according to the Wald-test statistic (Supplementary Data 4). Pathway annotation for *C. rodentium* was retrieved from the Kyoto Encyclopedia of Genes and Genomes database (KEGG; https://www.genome.jp/kegg/pathway.html). Analysis was performed by using 100,000 gene set permutations using gene sets limits of 5-500. Only pathways that passed an FDR < 0.05 were considered as significantly enriched. Module over-representation analysis was performed using a separate pre-ranked list for up- and down-regulated genes against the background universe of all detected genes (after pre-filtering) using ClusterProfiler^46^ (v4.6.0) and a Benjamini-Hochberg (BH) adjusted *p*-value threshold of < 0.05. Enrichment analysis data are available in Supplementary Data 5-6.

### RT-qPCR analysis

*C. rodentium* was grown in 6-well plates with 3 mL of DMEM supplemented with either DMSO or 10 µM of C4-HSL for 6 hours at 37 °C with 5 % CO2. RNA was isolated by using the Qiagen RNeasy Mini Kit; gDNA was depleted using the Turbo DNA-free kit (Invitrogen, AM1907). Reverse Transcription was performed on 1 µg of RNA using a QuantiTect Reverse Transcription Kit (Qiagen), and qPCR was performed using a QuantiNova SYBR Green PCR kit (Qiagen). Samples were analyzed in a ViiA 7 Real-Time PCR System (Applied Biosystems). All primer pairs used had an efficiency between 98-103 % (data not shown); sequences are reported in Supplemental Table S5. Normalized fold expression was calculated by using the comparative CT method^47^ with *dnaQ* and *recA* as endogenous controls.

### Type III secretion assay

Analysis of type III secreted proteins was performed as previously described^25^. In brief, *C. rodentium* strains and mutants were diluted in DMEM from an overnight culture in LB and grown statically for 6 hours at 37 °C with 5 % CO2. Proteins secreted into the culture supernatant were precipitated using trichloroacetic acid, separated by a 16 % SDS-PAGE gel, and stained using Coomassie Blue G250.

### Bacterial attachment to CMT-93 cells

Mouse intestinal epithelial cells (CMT-93; ATCC CCL-223) were grown and maintained in DMEM+++ at 37 °C and 5 % CO2. Before infection, cells were grown to confluence in a 12-well plate. A 3.5-hour subculture of WT *C. rodentium*, deletion mutants or complemented strains grown in LB to mid-log phase were used to infect cells to a multiplicity of infection (MOI) of 100. Upon inoculation, plates were spun for 5 minutes at 1000 rpm to synchronize the infection. At 4 hours post infection, cells were washed 5 times with PBS −/− to remove any non-adherent bacteria. Epithelial cells were lysed using 0.1 % Triton X-100, and tissue adherent bacteria were quantified via CFU plating.

### Cytometric Bead Array (CBA) assay

At days 4 and 7 post-murine infection, cecal and colonic tissues were homogenized in 1 mL of PBS −/− supplemented with the cOmplete EDTA-free protease inhibitor cocktail (Roche Diagnostics). Supernatants were collected by centrifuging the homogenate at 16,000 ×*g* at 4 °C for 20 minutes. Cytokine concentrations were determined using a Mouse Inflammation Cytometric Bead Array Kit (CBA; BD Biosciences) as per the manufacturer’s instructions. Samples were analyzed on an Attune NxT Flow Cytometer (ThermoFisher Scientific). Cytokine levels were normalized according to the sample weight.

### Data and statistical analysis

Statistical analyses were performed using Graphpad Prism (v9.3.1) (www.graphpad.com) or R (v4.1.0). Data were reported as the mean ± SD unless otherwise indicated. Sample size or number of replicates are indicated in the figure captions. When appropriate, significance was assessed by using a t-test or an ANOVA; for non-normal distributed data a Kruskal–Wallis or Mann-Whitney test was used. Post-tests are reported in the figure captions. Statistical significance is reported as follows: \**p* ≤ 0.05; \*\**p* < 0.01; \*\*\**p* < 0.001; \*\*\*\**p* < 0.0001.

## Results

### *C. rodentium* synthesizes multiple AHLs when grown *in vitro*

To determine the role that AHLs have in the regulation of *C. rodentium* behaviour, we first sought to profile the pathogen’s AHL production profile across time. To detect and quantify autoinducers present in complex mixtures, a LC-MS/MS method was developed. In brief, cell-free supernatants were obtained by growing *C. rodentium* in various growth media (YCFAG, DMEM, DMEM+++, and LB) across various timepoints (6-, 8-, 12- and 24-hours post-inoculation). AHLs were partially purified by using ethyl-acetate, and extracts were analysed via LC-MS/MS. Our results showed that *C. rodentium* primarily synthesizes C4-HSL as the dominant AHL signal with a peak average concentration of 7,973.4 nM observed at 12 hours post-inoculation when grown in YCFAG. Additionally, C6-HSL and 3-hydroxy-C6-HSL were also detected at lower concentrations, with peak average levels of 21.8 nM and 0.3 nM at 12 hours post-inoculation respectively (Figure 1A and Figure S2). Contrary to previous suggestions hinting that *C. rodentium* also synthesizes 3-oxo-C6-HSL^48^, there was no detection of 3-oxo-C6-HSL in any of the conditions tested. Our results also showed, as with many other QS systems, that the peak concentration for all detected AHLs was attained near the stationary phase (12 hours post-inoculation). However, a sharp reduction in all detected AHLs was observed at the stationary phase (24 hours post-inoculation), suggesting the presence of an uncharacterized regulatory mechanism in *C. rodentium* that may control the production or degradation of AHLs at high bacterial cell densities^49^.

**Figure 1:**
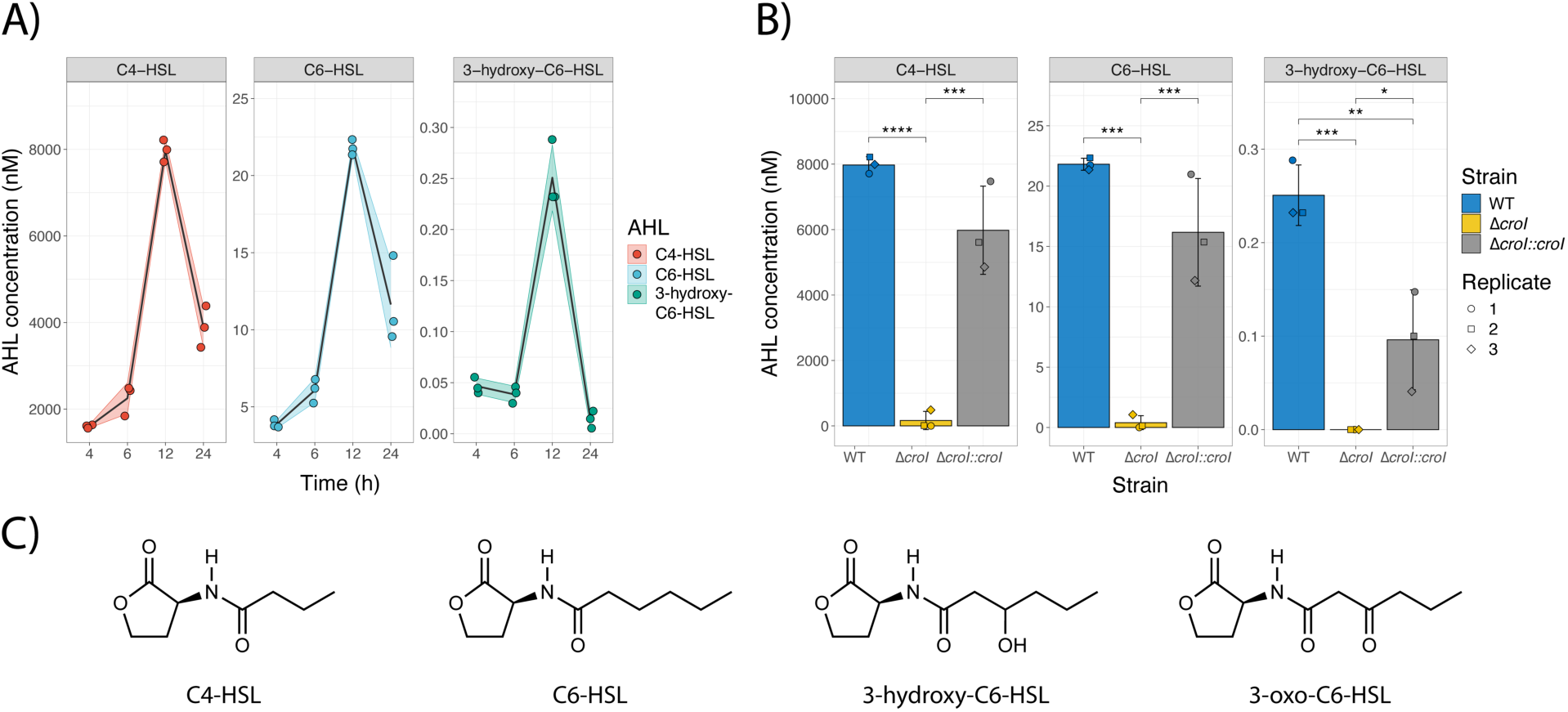
AHL production during *Citrobacter rodentium* growth. **A)** AHL production profile of WT *C. rodentium* grown over time in YCFAG (Yeast extract, Casitone and Fatty Acid Glucose media) as analyzed via LC-MS/MS. **B)** Comparison of AHL production by WT *C. rodentium*, the Δ*croI* mutant, and the complemented strain analyzed at 12 hours post-inoculation in YCFAG media. The values reported for the WT strain are the same as those reported in panel A at 12 hours post-inoculation. **C)** Chemical structures of the different N-acyl-homoserine lactones (AHLs) tested: N-butyryl-L-homoserine lactone (C4-HSL), N-hexanoyl-L-homoserine lactone (C6-HSL), N-(β-ketocaproyl)-L-homoserine lactone (3-oxo-C6-HSL), 3-hydroxy-hexanoyl-L-homoserine lactone (3-hydroxy-C6-HSL). **A)** Black line represents the mean values across time, shaded areas represent the standard deviation. **B)** Data are represented as the mean and standard deviation of three independent replicates. Significance was determined by a one-way ANOVA with Tukey’s for multiple comparisons test.

To test whether CroI, the only predicted AHL synthase encoded in *C. rodentium*, is responsible for synthesizing the multiple AHLs detected, an AHL-synthase deletion mutant (Δ*croI*) and its complemented strain were created. Deletion of *croI* resulted in a significant reduction of all detected AHLs, whereas chromosomal complementation of the AHL synthase (Δ*croI::croI*) resulted in the restoration of AHL synthesis to levels resembling those by the WT strain (Figure 1B). These data demonstrate that *C. rodentium* utilizes the AHL-synthase CroI to synthesize C4-HSL, C6-HSL and 3-hydroxy-C6-HSL (chemical structures in Figure 1C).

### RNA-seq revealed that AHL-mediated QS regulates virulence and metabolism pathways in *C. rodentium*

To gain insight into which genes are regulated by the AHL-mediated QS system of *C. rodentium*, we performed RNA-seq comparing the gene expression profile between the wild-type (WT) and Δ*croI* strains. We identified that in the absence of AHL production, there were 92 transcripts differentially upregulated and 125 transcripts differentially downregulated (Figure 2A; differential expression data are available in Supplementary Data 3). Gene set enrichment analysis (GSEA) identified an enrichment in pathways related to bacterial secretion systems and to flagellar assembly; as well as a downregulation of genes associated with starch and sucrose metabolism (Figure 2B, data available in Supplementary Data 5). KEGG module over-representation analysis further revealed that differentially upregulated genes in the Δ*croI* strain were strongly associated specifically with the type III secretion (T3S) system and associated effectors (Figure 2B, Supplementary Data 6). Indeed, we found that 35 out of the 41 LEE-encoded genes were significantly upregulated in the Δ*croI* strain, including several genes encoding transcriptional regulators and key structural, chaperone, translocator, and effector components of the T3SS (Figure 2C), as well as other non-LEE encoded T3S-related effector genes (*espT, espN2-2, espJ, espS,* and *espN1*). In contrast, some of the most significantly downregulated genes included genes associated with stress response (*rpoS*, *osmY*, *elaB* and *katE*), and with starch and sucrose metabolism (*bcsA, treZ, treY, treF, otsA, amyA, glgX* and *glgC*). These results are significant given the established importance that the interplay between stress-response, metabolism and virulence have in the successful colonization of a host by a pathogen^50, 51^. Together, these results indicate that *C. rodentium* is capable of detecting changes in AHL levels in the environment and using them as regulatory signals to initiate targeted changes in gene expression involved in virulence, stress and metabolism.

**Figure 2:**
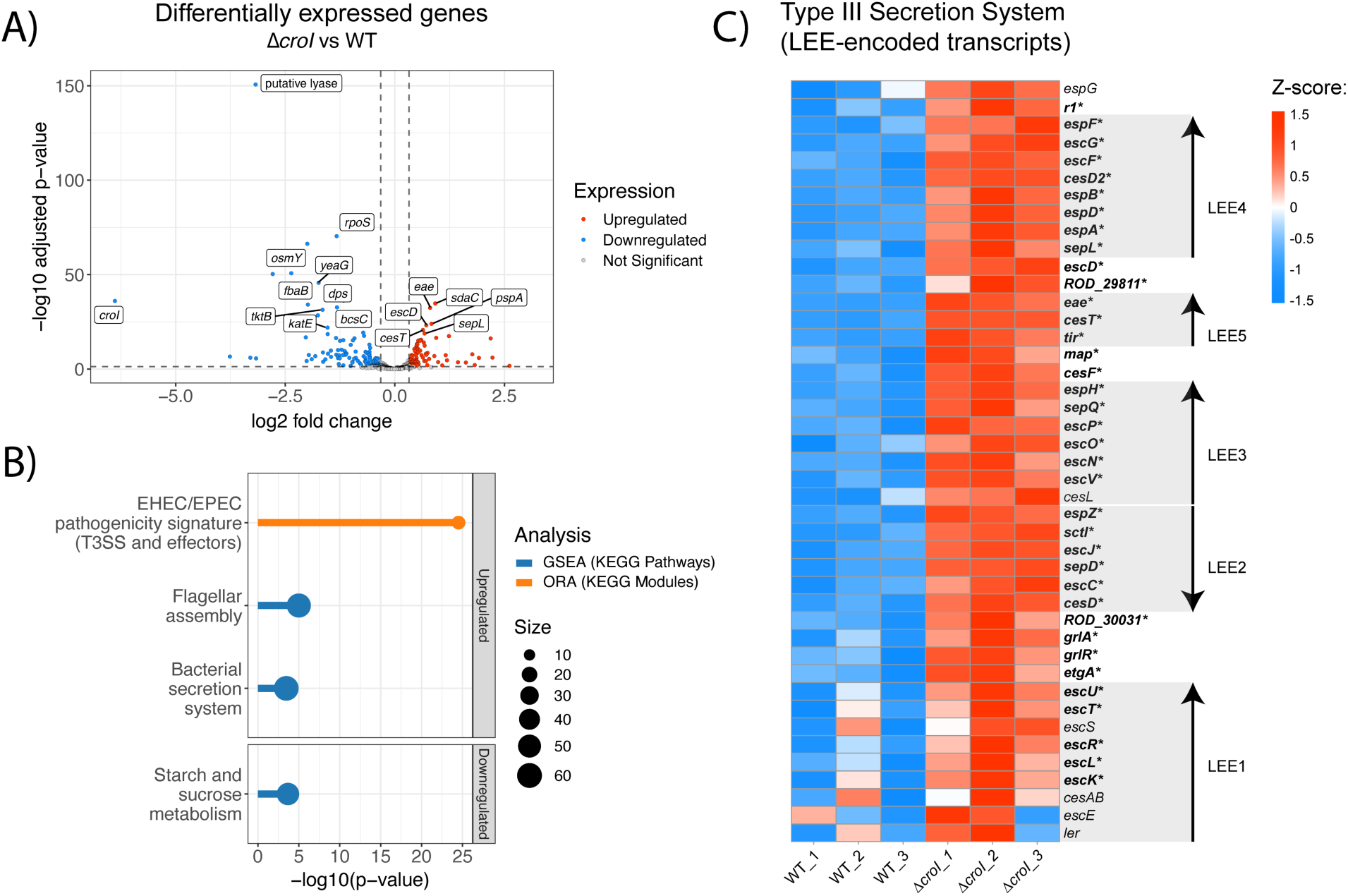
QS regulates expression of the type III secretion system in *Citrobacter rodentium*. **A)** Volcano plot showing differentially expressed genes (DEGs) from RNA-Seq data comparing the Δ*croI* strain to WT *C. rodentium*. Genes shown in red are significantly upregulated with a fold change greater than 1.25 and *p*-value *<* 0.05, genes in blue are significantly downregulated with a fold change lower than 1.25 and *p*-value *<* 0.05. The top 20 significantly differentially expressed genes ordered by ascending *p*-value are labelled. Differential expression data are available in Supplementary Data 3. **B)** Pathway enrichment analyses were calculated by using Gene Set Enrichment Analysis (GSEA; shown in blue); and overrepresented KEGG modules (shown in orange) were calculated by using ClusterProfiler. Only pathways that passed an FDR *<* 0.05 were considered significantly enriched. Supplementary enrichment data are available in Supplementary Data 5-6. **C)** Heatmap showing Z-score expression of genes associated with the type III secretion system. Each operon within the Locus of Enterocyte Effacement (LEE1-5) is displayed using arrowheads to indicate gene orientations^63^. Significantly differentially regulated genes when comparing Δ*croI* and WT are shown as bolded text with an asterisk at the end of the gene name. ORA, over-representation analysis.

### QS regulates T3SS activity and epithelial cell attachment of *C. rodentium*

To confirm whether the observed increase in T3S-associated transcript levels corresponds to an increase in translation and secretion of type III-associated proteins, a T3S assay was performed. The Δ*croI* strain displayed a 49 % average increase in the abundance of secreted proteins as compared to WT *C. rodentium* (Figure 3A), while chromosomal complementation of the AHL synthase (Δ*croI::croI*) restored the protein secretion profile back to WT levels. To further disentangle the AHL-mediated CroI/CroR signaling system, we then asked whether deletion of the AHL-receptor CroR would also induce an increase in secretion of type III-associated proteins. Interestingly, Δ*croR* and its complemented strain (Δ*croR::croR*) showed no changes in their type III secretion profile when compared to the WT strain. This suggests that CroR behaves as a transcriptional repressor, whereby changes in the expression of QS-regulated genes can only occur after transcriptional repression is relieved following the binding of an AHL to its cognate AHL-receptor. This observation is consistent with a previous study by Colthurst, et al.^48^, where they show no phenotypical changes between WT and a Δ*croR* strain. These data confirm that QS has a significant role in the regulation of the T3SS, and likely contributes to an enhancement of virulence under conditions with low bacterial densities where AHL levels are minimal.

**Figure 3:**
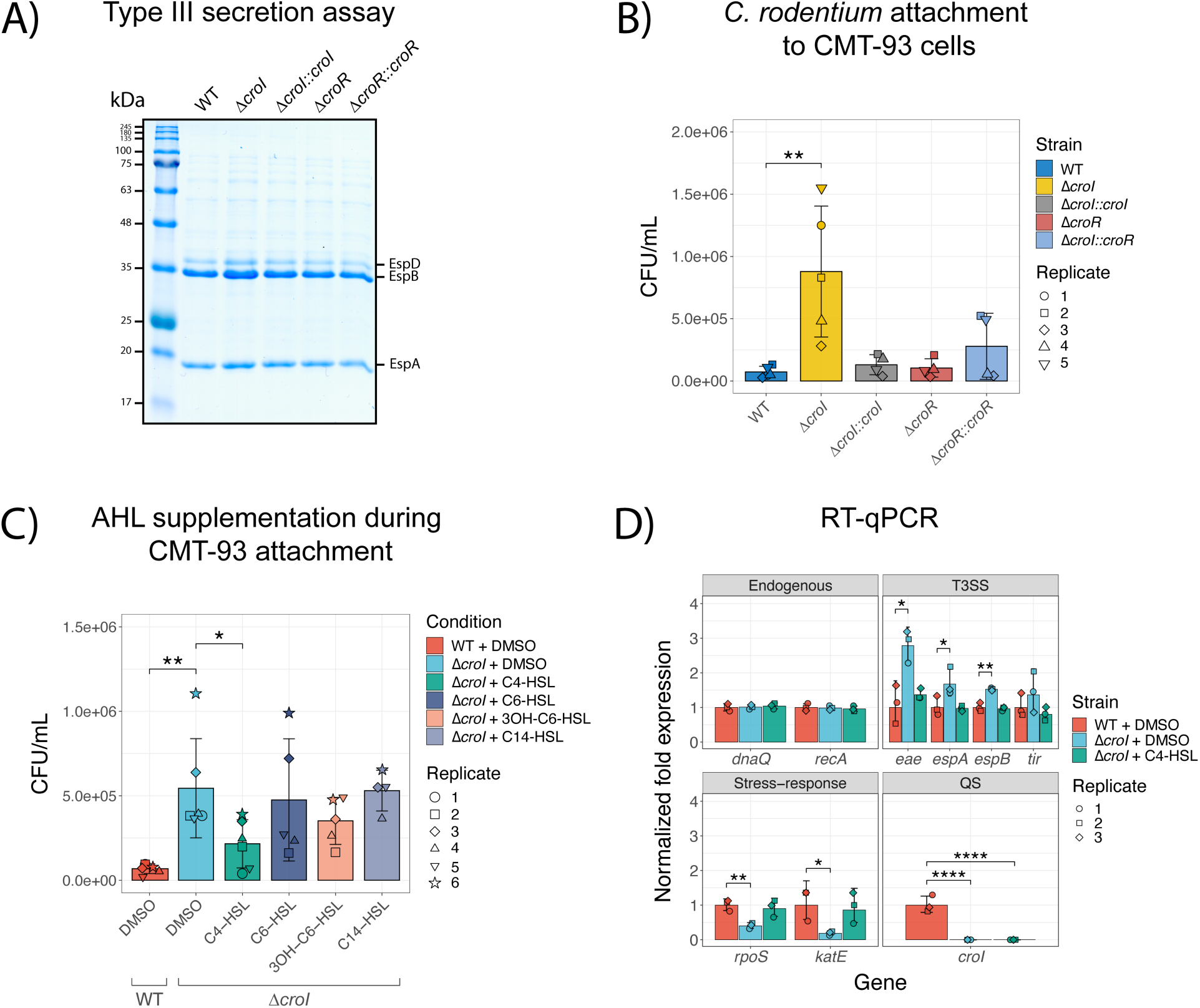
CroI plays a role in regulation of Type III Secretion (T3S) and bacterial attachment to host epithelial cell surfaces. **A)** Bacterial attachment of WT *C. rodentium*, Δ*croI*, Δ*croR*, and complemented strains to CMT-93 murine epithelial cells 4 hours post-infection. **B)** Attachment of WT and Δ*croI* strains to CMT-93 cells in response to supplementation of culture media with various AHLs (10 µM) or DMSO vehicle control. Data are represented as the mean and SD of at least four independent replicates (N=4-6). Significance was determined using pairwise t-tests with Bonferroni corrected *p*-values for multiple comparisons. **C)** Profile of T3SS secreted proteins derived from the supernatants of WT, Δ*croI*, Δ*croR*, and complemented strains as shown on an SDS-16 % PAGE gel stained with Coomassie Blue G250. The Δ*croI* strain displayed a 49 % average increase in the abundance of secreted proteins as compared to WT *C. rodentium* (average measured across 3 different gels). **D)** RT-qPCR gene expression analysis of selected differentially expressed genes identified through RNA sequencing. *dnaQ* and *recA* were used as endogenous controls. All primers had an efficiency between 98-103 %. Results are the average of 3 technical replicates prepared from strains grown in DMEM in biological triplicate cultures. Bar heights and error bars represent geometric means ± standard deviation. Statistics were calculated by using multiple t-tests with Bonferroni correction for multiple comparisons. T3SS, Type III Secretion System; QS, Quorum Sensing; 3OH-C6-HSL, 3-hydroxy-C6-HSL.

Given the observed upregulation in T3SS activity, we examined whether disruption of CroI could lead to additional functional enhancements in *C. rodentium* virulence responses during infection. One of the hallmark characteristics of A/E infection is the intimate attachment to the host epithelium, and so *in vitro* assays were designed to assess bacterial adherence to intestinal epithelial cells, with the aim of modelling conditions within the murine gut. To achieve this, a mouse rectal carcinoma cell line (CMT-93 cells) was utilized. Indeed, the efficiency of *C. rodentium* attachment to CMT-93 cells was significantly increased in the *ΔcroI* mutant (Figure 3B). Consistent with our previous results, we did not identify any differences in epithelial cell attachment for the Δ*croR* and Δ*croR*::*croR* strains when compared to WT. If the absence of AHLs in the environment enhances epithelial adherence, we hypothesized that exogenous supplementation of AHLs to the non-secreting Δ*croI* strain would chemically complement the CMT-93 attachment phenotype. In support of this hypothesis, we found that indeed C4-HSL was capable of consistently reverting this phenotype by restoring Δ*croI* epithelial attachment to levels similar to WT; while no significant differences were observed in any other of the tested AHLs (Figure 3C). This phenotype was not due to any changes in *C. rodentium* growth dynamics caused by the supplementation of AHLs into the culture media (Figure S3). Collectively, these findings show that in the absence of AHLs, *C. rodentium* can increase bacterial attachment to host epithelial cells; additionally, they demonstrate that C4-HSL plays a crucial role as a signaling molecule directly influencing the virulence response of *C. rodentium*.

To verify the modulatory effects of C4-HSL, we measured changes in expression of key genes identified from our RNA-seq screen by using RT-qPCR. As expected, we confirmed that several T3SS-related genes were significantly upregulated when AHL synthesis was disrupted; in contrast, stress-related genes were shown as significantly downregulated in the Δ*croI* strain when compared to WT (Figure 3D). Furthermore, we confirmed that exogenous addition of C4-HSL to the culture media chemically complemented Δ*croI* by reverting transcript expression to WT levels. Together with our functional analysis, these data suggest that the production and detection of C4-HSL plays a critical role in the regulation of *C. rodentium* virulence and bacterial stress responses, and may therefore have an essential role in pathogen adaptation during host colonization.

### AHLs can be detected across the gastrointestinal tract during a time course of infection

Having characterized the link between the AHL-mediated QS system of *C. rodentium* and pathogen behaviour *in vitro*, we next focused on investigating the role of QS during host infection. Studying *C. rodentium* offers the rare opportunity to directly profile the concentration of target AHLs present within the murine gut both in naïve mice, and over the time course of infection. Adult C57BL/6J mice were gavaged with either WT *C. rodentium, ΔcroI,* or with PBS as a control. AHLs were then partially purified from regional gut contents and extracts were analyzed via LC-MS/MS. To capture both early- and peak-infection stages, intestinal contents were collected at 4- and 7-days post-infection (p.i.), focusing specifically on the cecum and colon which are important sites of initial *C. rodentium* colonization (Figure 4A-B). We detected and quantified C4-HSL in WT colonized mice at both day 4 and 7 p.i., and in both the cecum and colon (Figure 4C-D; with an average of 0.10 and 12.80 ng/g in the cecum, and 3.71 and 21.50 ng/g in the colon, at day 4 and 7 p.i. respectively), confirming C4-HSL as the dominant AHL produced by *C. rodentium* during infection. Low concentrations of C6-HSL were also detected, primarily within the colon during peak infection (average of 0.06 and 0.72 ng/g at day 4 and 7 p.i. respectively). Despite demonstrating the capacity of *C. rodentium* to synthesize 3-hydroxy-C6-HSL, it could not be detected in any of the tested samples. Since detected 3-hydroxy-C6-HSL concentrations remained low even within saturated culture conditions (Figure 1A), it is likely that the accumulated levels of 3-hydroxy-C6-HSL within the gut contents are lower than our limit of detection. No 3-oxo-C6-HSL was detected in any of the samples tested. Both C4-HSL and C6-HSL were only detected in samples where *C. rodentium* reached a bacterial burden greater than ∼10^7^ CFUs, indicating that our methodology requires a minimum threshold of pathogen density in order to detect AHLs (Figure S4A and Figure S5C).

**Figure 4:**
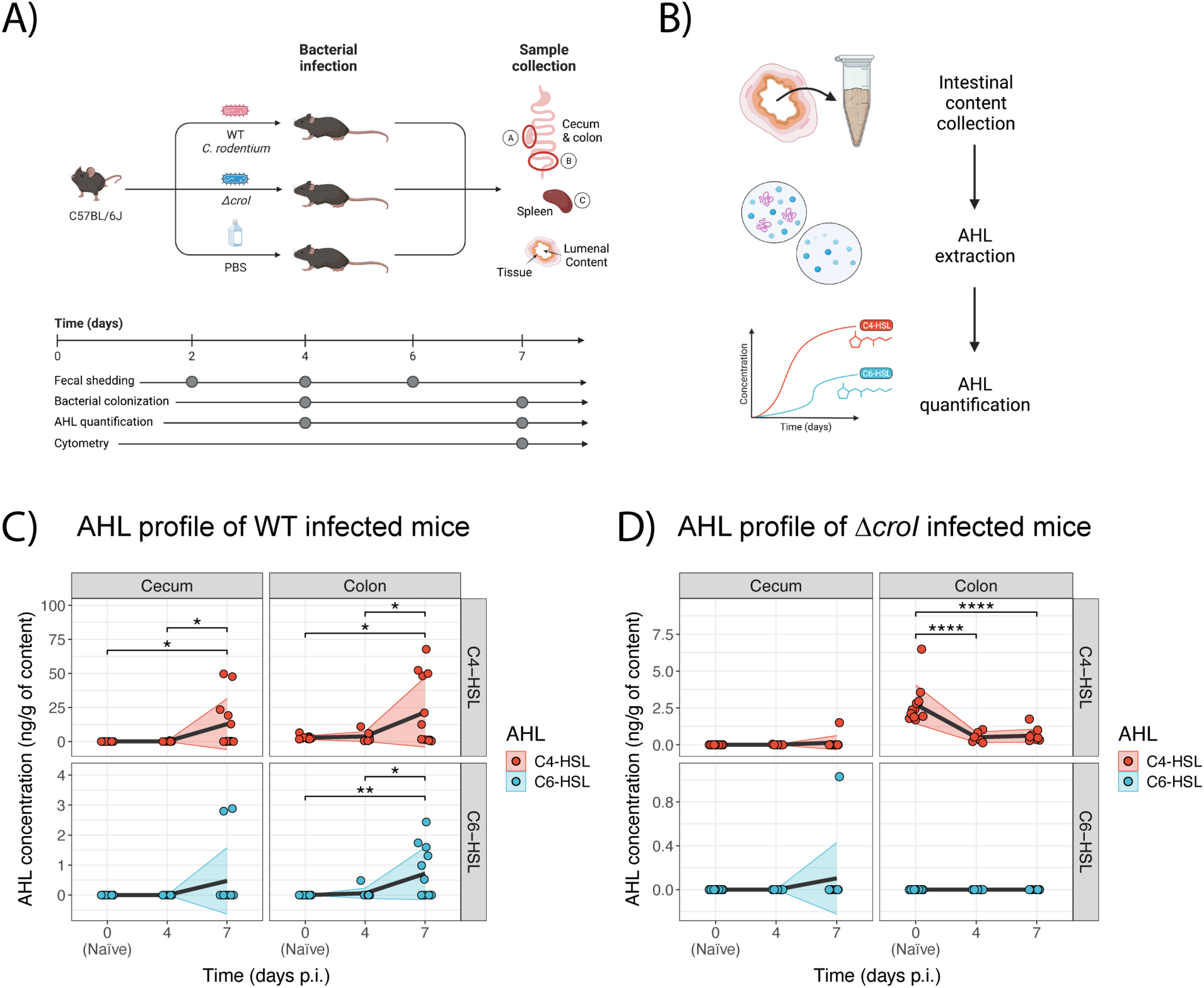
Multiple AHLs can be detected *in vivo* over the course of *C. rodentium* infection. **A)** Graphical representation of the murine infection set up. In brief, C57BL/6J mice were gavaged with either WT *C. rodentium*, Δ*croI*, or PBS (uninfected control group). At experimental endpoint large intestinal (cecum and colon) and systemic (spleen) organs were collected for CFU enumeration. **B)** Cecum and colon contents were also collected on days 4 and 7 post-infection (p.i.) for partial purification and detection of N-acyl-homoserine lactones (AHLs) via LC-MS/MS. **C-D)** Changes in concentration of C4-HSL and C6-HSL detected in cecum and colon contents during **C)** WT and **D)** Δ*croI* infection. 3-hydroxy-C6-HSL and 3-oxo-C6-HSL were not detected. As a reference, the values detected for the naïve mice were plotted on both panels C and D as day 0 post-infection. The black line represents the mean values across time, and shaded areas represent the standard deviation with an N =6-12 mice per group. Statistics were calculated by using multiple t-tests with Bonferroni correction for multiple comparisons.

To further investigate the potential role of AHLs during infection, we next correlated the regional *C. rodentium* burden of individual mice to their corresponding AHL concentrations. We found that both C4-HSL and C6-HSL levels were positively associated with both lumenal (Figure S4A; C4-HSL rho = 0.92, *p* < 0.001; C6-HSL rho = 0.86, *p* < 0.001), and tissue-associated (Figure S5C; C4-HSL rho = 0.65, *p* = 0.026; C6-HSL rho = 0.68, *p* < 0.014) *C. rodentium* subpopulations in the colon. This association was not observed in the cecum (Figure S4A and Figure S5C); this may be due to the high degree of variability associated with this intestinal site where factors such as food consumption (further diluting AHL concentrations) or the high variation in the consistency and amount of intestinal content available between different mice made the detection of cecal AHLs particularly challenging. Remarkably, low levels of C4-HSL were also detected in colonic samples of naïve mice (Shown as day 0 p.i. in Figure 4C-D; with an average of 2.74 ng/g) despite having no detectable colonization of *C. rodentium* (as measured via selective plating). These findings suggest that there might be members of the resident microbiota capable of synthesizing AHLs.

We next examined how AHL concentrations change after infection with the Δ*croI* mutant strain. We found that detected C4-HSL levels were significantly reduced in Δ*croI*-infected mice when compared to the naïve samples (Figure 4D; average of 0.51 ng/g and 0.63 ng/g detected in the colon at day 4 and 7 p.i. respectively). Furthermore, our data revealed that AHL levels tended to decrease with increasing *ΔcroI* burden (Figure S4B), further supporting the possibility that members of the resident microbiota can synthesize C4-HSL, and that the invading non-AHL producing Δ*croI* strain is displacing any AHL-producing commensal strains from their natural intestinal niche, leading to a loss of microbiota-derived C4-HSL from the gut. Altogether, these data validate the production of AHLs within the gut, not only by *C. rodentium* during infection, but possibly also by the resident microbiota. Furthermore, the temporal accumulation of *C. rodentium-*produced AHLs in *vivo*, suggests that the production and detection of AHLs is likely a key regulatory signal used by *C. rodentium* to coordinate infection stage-specific gene expression.

### *C. rodentium* utilizes AHLs to regulate colonization dynamics and control host damage during infection

To further expand on the role of the AHL-mediated QS system of *C. rodentium* during host infection, we assessed the colonization dynamics of both the WT and Δ*croI* strains following inoculation of C57BL/6 mice. Enumeration of *C. rodentium* shed in the feces daily post-infection revealed a significant increase in bacterial shedding over the course of Δ*croI* infection (Figure 5A; Mixed-effects model with Geisser-Greenhouse correction, Time × Strain: *p* = 0.0037), indicating a possible role for QS in pathogen transmissibility to new hosts. On days 4 and 7 post-infection, we assessed pathogen burden across GI tissues, as well as within the spleen to represent systemic spread. We found no significant differences in pathogen burden between the Δ*croI* and WT strains in neither lumen-associated *C. rodentium* subpopulations in the cecum or colon (Figure 5B-C), nor in tissue-associated subpopulations (Figure S5A-B). However, when measuring systemic burden, we observed that by day 7 p.i. the Δ*croI* strain had a 100 % rate of systemic spread, as compared to 50 % during WT infection (Figure 5D; Fisher’s exact test, *p* = 0.015); as well as an overall increase in spleen pathogen burden (Figure 5E). We believe that this is likely a consequence of the increased expression in virulence machinery from the Δ*croI* strain leading to an increase in damage to the epithelial cell barrier, consequently increasing translocation of bacteria entering host circulation. Analysis of the tissue-derived cytokine profile across the large intestine did not identify any major changes between WT- and Δ*croI*-infected tissues, although there was a trend towards decreased levels of cecal TNFα in the context of Δ*croI* infection (Figure S6). These data indicate that pathogen transmissibility and systemic spread may be influenced by QS, possibly as an important strategy to coordinate the expression of virulence genes during *in vivo* infection.

**Figure 5:**
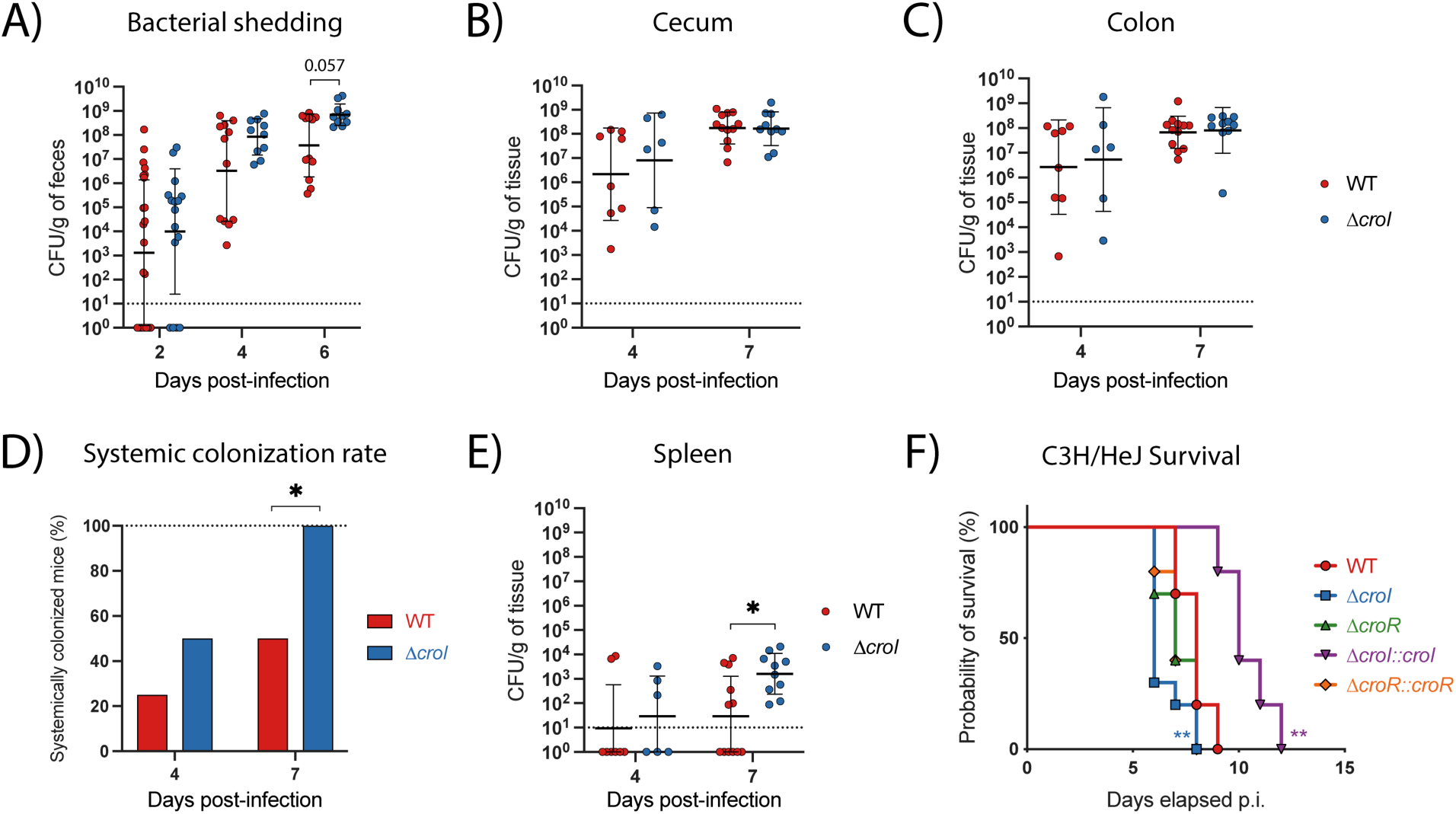
*C. rodentium* Δ*croI* displays increased pathogenicity in C57BL/6J and C3H/HeJ mouse infection models. **A)** Fecal shedding of *C. rodentium* across the infection time course. Data were analyzed using a Wilcoxon rank test using Holm-Šidák correction for multiple comparisons. **B-C)** *C. rodentium* burden in cecal and colonic contents measured at day 4 and 7 post-infection (p.i.). **D-E)** Systemic colonization found in whole spleens at day 4 and 7 post-infection. **D)** Success rate of *C. rodentium* spleen colonization. Statistical analysis was calculated using Fisher’s exact test. **E)** CFU burden found in whole spleens. **F)** Survival of susceptible C3H/HeJ mice infected with WT, Δ*croI,* Δ*croR* and complemented strains of *C. rodentium* reveals an increased morbidity and mortality in mice infected with the Δ*croI* strain as compared to WT infection. For WT and each of the mutant strains tested an N = 10 mice was used, and for complemented strains an N = 5 was used instead. Statistics were calculated using a logrank (Mantel-Cox) test comparing each strain against WT *C. rodentium.* **A-C, E)** Lines represent geometric mean ± geometric standard deviation, with an N = 6-12 mice per group. The limit of detection was displayed as a dotted line. Statistical analysis was performed by using a Mann-Whitney test.

Given the observed increase in systemic spread in mice infected with the Δ*croI* strain, we sought to determine whether interfering with AHL synthesis altered host damage or displayed any changes to the lethality of infection. This was done by infecting C3H/HeJ mice, which are highly susceptible to *C. rodentium* and succumb to 100 % lethality. Indeed, mice gavaged with the Δ*croI* strain displayed hypervirulence, resulting in an increase in morbidity and mortality of the mice as compared to WT infection (Figure 5F). In contrast, this phenotype was successfully reversed upon restoration of AHL-production during infection with the Δ*croI::croI* complemented strain. Deletion of the AHL-receptor CroR or its complemented strain (Δ*croR::croR)* did not result in differences to host disease outcome as compared to WT infection, once again supporting its role as a negative regulator. Overall, these data indicate that interference with AHL-mediated QS systems in *C. rodentium* results in increased pathogenesis and host disease severity.

## Discussion

QS dynamics within the mammalian gastrointestinal tract remains poorly understood, and many questions remain regarding the composition and abundance of different AHLs across distinct GI regions, the kinetics of AHL synthesis and degradation, and the potential regulatory capacity that AHLs have as signaling molecules for both commensal and pathogenic microbes. In this study, we show that AHLs can be detected across different regions of the mammalian gastrointestinal tract both during health and disease, hinting towards their use by both pathogens and commensals during niche colonization. We also highlight the regulatory capacity that AHLs have during a natural model of infection by demonstrating how *C. rodentium* utilizes QS-mediated signaling to control the timing of expression of virulence genes to maximize its chances to successfully colonize its host.

The detection of AHLs from the mammalian intestine has been notoriously elusive. Reasons for this include low accumulation of AHLs found *in vivo*, low sensitivity and biases associated with the use of biosensors, difficulty to detect a wide array of molecular structures of AHLs, and possible degradation of AHLs by either the host or the microbiota^17^. To date, only a handful of studies have detected AHLs from *in vivo* samples. These observations have mostly been indirect, either by using biosensors^52^, or by using mass spectrometry to analyze fecal samples derived from IBD patients^53^. To our knowledge, only one study has successfully detected AHLs directly from biological samples by analyzing cecal contents, sera and liver of mice^54^. Our study not only supports previous findings by showing that AHLs can be detected *in vivo,* but further demonstrates how AHL composition changes across multiple intestinal sites throughout the course of infection (Figure 4). While we observed similar abundance of C4-HSL in cecal contents of infected animals to Xue et al.^54^, we were also able to detect low levels of C6-HSL, a secondary AHL molecule synthesized by *C. rodentium* (Figure 4C). An interesting observation was the overall increase in AHL levels in colonic contents when compared with cecal contents in both infected and uninfected animals. These findings suggest the existence of ecological forces affecting QS dynamics within the gastrointestinal tract, such as the effects of complex topology present within intestinal colonic crypts leading to the prevention of AHL flowthrough^55^, or the effects of long-range flow from bacteria colonizing upstream in the intestine allowing for higher concentrations of AHLs to accumulate downstream^56^.

It is rare to be able to investigate the role of a QS system during pathogen infection of its natural host, and studying *C. rodentium* offered us the opportunity to characterize the AHL-mediated QS regulon both *in vitro* and *in vivo.* By studying a strain lacking an AHL-synthase (Δ*croI*), we were able to recreate conditions which mimic the environment that *C. rodentium* encounters as it initially enters the gastrointestinal tract (when AHL concentrations are likely minimal). Our results showed that in the absence of AHLs, *C. rodentium* promotes the expression of the T3SS (Figure 3), highlighting the role that the absence of AHLs has as a regulatory cue stimulating the pathogen to prepare for its intimate attachment to the host epithelia. This relationship is likely context dependent, as previous studies have shown that a Δ*croI* mutant strain had reduced attachment to non-biotic surfaces^48^. Collectively, this shows that QS regulation of bacterial behaviour according to its spatio-temporal context is critical for pathogen success, from coordinating behaviours that promote colonization during infection, such as increasing virulence machinery (like the T3SS) to allow for intimate attachment to the host epithelium, to those that lead to host damage, potential immune avoidance and even transmission to a new host (Figure 5 and Figure S6).

The fact that one synthase, CroI, could lead to the production of three distinct AHLs (Figure 1), supports the idea of heterogeneity in AI accumulation that can happen under a natural model of infection. This highlights possible mechanisms of host-adaptation that can arise from having subpopulations of isogenic bacteria existing in different phenotypical states which consequently lead to an overall increase in bacterial fitness of the population as a whole^2, 57^. Considering this, high accumulation of C4-HSL could be considered a late-stage infection signal indicating to the attached subpopulation of *C. rodentium* that they were successful in reaching their colonization niche and on establishing an infection, consequently triggering a downregulation in expression of the metabolically expensive T3S machinery. Alternately, high C4-HSL concentrations could also indicate the presence of another competing pathogen or microbiota member within the *C. rodentium* colonization niche, indicating that colonization conditions are not optimal and the expression of the T3SS could come at a disadvantage^58^. It would be interesting to explore what role C6-HSL and 3-hydroxy-C6-HSL have as signaling molecules during *C. rodentium* infection. One hypothesis is that they allow *C. rodentium* to expand its QS signaling network^59^ allowing for a fine-tuned regulation of pathogen behaviour according to the infection stage and geographic location within the gut. Alternately, they could potentially play a role as inter-species signaling molecules which could increase *C. rodentium*’s ability to outcompete other microbes that share its colonization niche^58^; or they could act as trans-kingdom signals capable of modulating host immune responses to increase *C. rodentium* colonization success rate^19, 20^.

Interfering with the QS communication of bacteria could provide a powerful strategy to control virulence of pathogenic bacteria. Strategies such as degradation or interference of the binding of an autoinducer to its cognate receptor can lead to repression of virulence thus allowing the immune system to naturally clear the infection without the need for additional interventions such as antibiotics^60, 61^. Results presented here, as well as by others, show that in the absence of AHLs some pathogens can display an increase in pathogenesis and host disease severity^10, 62–64^, highlighting that QS regulation must be understood in the context of infection of individual pathogens, and that development of future therapeutics targeting QS signaling needs to be pathogen specific, and should not be considered a generalized strategy.

In summary, our work demonstrates that AHLs can be detected in different regions from the mammalian gastrointestinal tract. These results have the potential to be expanded to other experimental systems and aid the understanding on how AHLs fluctuate throughout the gastrointestinal tract both in health and disease. Additionally, we demonstrated the ability of the enteric pathogen *C. rodentium* to utilize its QS system to produce, release and respond to AHLs to regulate its virulence and maximize colonization success. Through comprehensive analysis of AHL-mediated QS, we characterized both the temporal production of AHLs by *C. rodentium*, as well as their effect on gene expression, virulence behaviour, and host infection dynamics. Together, these findings demonstrate the important role that QS plays within the mammalian gut and further expand our understanding of QS regulation by gastrointestinal pathogens.

## Acknowledgements

The authors would like to thank all of our colleagues in the Finlay laboratory for their support and assistance. This work was supported by grants from the Canadian Institutes of Health Research (CIHR) to BB Finlay (FDN-159935). J Peña-Díaz received support from the University of British Columbia (UBC) and from Mitacs. Supporting images were created with BioRender (<u>BioRender.com</u>). This research was enabled in part by software provided by the Digital Research Alliance of Canada (<u>alliancecan.ca</u>). We are grateful to Matthew Croxen and Kirsten Koymans for generating the DBS100 Δ*croI* mutant. We would also like to thank Lisa Thorson for the fundamental logistical support of the project. We thank Christopher Lee and Zakhar Krekhno for the invaluable discussions on RNA-seq analysis.

## Author Contributions

J.P.D., and B.B.F conceived the project. J.P.D., S.E.W., A.S.P., A.C.C and W.D. performed and designed experiments. J.P.D., S.E.W., A.C.C., analyzed data. J.P.D. wrote bioinformatics pipelines. J.P.D. wrote the original draft of the manuscript with input from all authors. All authors revised the manuscript. B.B.F. acquired funding for the project and provided supervision.

## Data Availability

The RNA-seq data generated from this study will be made available on the NCBI Sequencing Read Archive (SRA) upon the final publication of the manuscript. Supplementary datasets will also be made available upon the final publication of the manuscript.

## References

1. Vogt, S.L., Peña-Díaz, J., and Finlay, B.B. (2015). Chemical communication in the gut: Effects of microbiota-generated metabolites on gastrointestinal bacterial pathogens. Anaerobe 34, 106–115. 10.1016/j.anaerobe.2015.05.002.

2. Woodward, S.E., Krekhno, Z., and Finlay, B.B. (2019). Here, there, and everywhere: How pathogenic Escherichia coli sense and respond to gastrointestinal biogeography. Cellular Microbiology 21, 1–15. 10.1111/cmi.13107.

3. Storz, G., and Hengge, R. (2010). Bacterial Stress Responses (American Society for Microbiology Press).

4. Miller, M.B., and Bassler, B.L. (2001). Quorum sensing in bacteria. Annu Rev Microbiol 55, 165–199. 10.1146/annurev.micro.55.1.165.

5. Waters, C.M., and Bassler, B.L. (2005). Quorum sensing: cell-to-cell communication in bacteria. Annual review of cell and developmental biology 21, 319–346. 10.1146/annurev.cellbio.21.012704.131001.

6. Rutherford, S.T., and Bassler, B.L. (2012). Bacterial quorum sensing: its role in virulence and possibilities for its control. Cold Spring Harbor perspectives in medicine 2, 1–26. 10.1101/cshperspect.a012427.

7. Nealson, K.H., and Hastings, J.W. (1979). Bacterial bioluminescence: its control and ecological significance. Microbiol Rev 43, 496–518.

8. Engebrecht, J., Nealson, K., and Silverman, M. (1983). Bacterial bioluminescence: Isolation and genetic analysis of functions from Vibrio fischeri. Cell 32, 773–781. 10.1016/0092-8674(83)90063-6.

9. de Kievit, T.R., and Iglewski, B.H. (2000). Bacterial Quorum Sensing in Pathogenic Relationships. Infect Immun 68, 4839–4849. 10.1128/IAI.68.9.4839-4849.2000.

10. Bronesky, D., Wu, Z., Marzi, S., Walter, P., Geissmann, T., Moreau, K., Vandenesch, F., Caldelari, I., and Romby, P. (2016). *Staphylococcus aureus* RNAIII and Its Regulon Link Quorum Sensing, Stress Responses, Metabolic Adaptation, and Regulation of Virulence Gene Expression. Annu. Rev. Microbiol. 70, 299–316. 10.1146/annurev-micro-102215-095708.

11. Davies, D.G., Parsek, M.R., Pearson, J.P., Iglewski, B.H., Costerton, J.W., and Greenberg, E.P. (1998). The Involvement of Cell-to-Cell Signals in the Development of a Bacterial Biofilm. Science 280, 295–298. 10.1126/science.280.5361.295.

12. Hammer, B.K., and Bassler, B.L. (2003). Quorum sensing controls biofilm formation in Vibrio cholerae: Biofilms in V. cholerae. Molecular Microbiology 50, 101–104. 10.1046/j.1365-2958.2003.03688.x.

13. Kong, K.-F., Vuong, C., and Otto, M. (2006). Staphylococcus quorum sensing in biofilm formation and infection. International Journal of Medical Microbiology 296, 133–139. 10.1016/j.ijmm.2006.01.042.

14. Fuqua, C., Parsek, M.R., and Greenberg, E.P. (2001). Regulation of Gene Expression by Cell-to-Cell Communication: Acyl-Homoserine Lactone Quorum Sensing. Annu. Rev. Genet. 35, 439–468. 10.1146/annurev.genet.35.102401.090913.

15. Papenfort, K., and Bassler, B.L. (2016). Quorum sensing signal–response systems in Gram-negative bacteria. Nature Reviews Microbiology 14, 576–588. 10.1038/nrmicro.2016.89.

16. von Bodman, S.B., Willey, J.M., and Diggle, S.P. (2008). Cell-Cell Communication in Bacteria: United We Stand. J Bacteriol 190, 4377–4391. 10.1128/JB.00486-08.

17. Swearingen, M.C., Sabag-Daigle, A., and Ahmer, B.M.M. (2013). Are There Acyl-Homoserine Lactones within Mammalian Intestines? J Bacteriol 195, 173–179. 10.1128/JB.01341-12.

18. Thompson, J.A., Oliveira, R., Djukovic, A., Ubeda, C., and Xavier, K.B. (2015). Manipulation of the quorum sensing signal AI-2 affects the antibiotic-treated gut microbiota. Cell Reports 10, 1861–1871. 10.1016/j.celrep.2015.02.049.

19. Coquant, G., Grill, J.-P., and Seksik, P. (2020). Impact of N-Acyl-Homoserine Lactones, Quorum Sensing Molecules, on Gut Immunity. Front. Immunol. 11, 1827. 10.3389/fimmu.2020.01827.

20. Coquant, G., Aguanno, D., Brot, L., Belloir, C., Delugeard, J., Roger, N., Pham, H.-P., Briand, L., Moreau, M., de Sordi, L., et al. (2022). 3-oxo-C12:2-HSL, quorum sensing molecule from human intestinal microbiota, inhibits pro-inflammatory pathways in immune cells via bitter taste receptors. Sci Rep 12, 9440. 10.1038/s41598-022-13451-3.

21. Hsiao, A., Ahmed, A.M.S., Subramanian, S., Griffin, N.W., Drewry, L.L., Petri, W.A., Haque, R., Ahmed, T., and Gordon, J.I. (2014). Members of the human gut microbiota involved in recovery from Vibrio cholerae infection. Nature 515, 423–426. 10.1038/nature13738.

22. Collins, J.W., Keeney, K.M., Crepin, V.F., Rathinam, V.A.K., Fitzgerald, K.A., Finlay, B.B., and Frankel, G. (2014). Citrobacter rodentium: Infection, inflammation and the microbiota. Nature Reviews Microbiology 12, 612–623. 10.1038/nrmicro3315.

23. Crepin, V.F., Collins, J.W., Habibzay, M., and Frankel, G. (2016). Citrobacter rodentium mouse model of bacterial infection. Nat Protoc 11, 1851–1876. 10.1038/nprot.2016.100.

24. Schauer, D.B., and Falkow, S. (1993). Attaching and effacing locus of a Citrobacter freundii biotype that causes transmissible murine colonic hyperplasia. Infect Immun 61, 2486–2492. 10.1128/iai.61.6.2486-2492.1993.

25. Deng, W., Vallance, B.A., Li, Y., Puente, J.L., and Finlay, B.B. (2003). Citrobacter rodentium translocated intimin receptor (Tir) is an essential virulence factor needed for actin condensation, intestinal colonization and colonic hyperplasia in mice. Molecular Microbiology 48, 95–115. 10.1046/j.1365-2958.2003.03429.x.

26. McDaniel, T.K., Jarvis, K.G., Donnenberg, M.S., and Kaper, J.B. (1995). A genetic locus of enterocyte effacement conserved among diverse enterobacterial pathogens. Proc. Natl. Acad. Sci. U.S.A. 92, 1664–1668. 10.1073/pnas.92.5.1664.

27. Deng, W., Li, Y., Vallance, B.A., and Finlay, B.B. (2001). Locus of Enterocyte Effacement from *Citrobacter rodentium*: Sequence Analysis and Evidence for Horizontal Transfer among Attaching and Effacing Pathogens. Infect Immun 69, 6323–6335. 10.1128/IAI.69.10.6323-6335.2001.

28. Deng, W., Puente, J.L., Gruenheid, S., Li, Y., Vallance, B.A., Vázquez, A., Barba, J., Ibarra, J.A., O’Donnell, P., Metalnikov, P., et al. (2004). Dissecting virulence: Systematic and functional analyses of a pathogenicity island. Proc. Natl. Acad. Sci. U.S.A. 101, 3597–3602. 10.1073/pnas.0400326101.

29. Gibson, D.G., Young, L., Chuang, R.-Y., Venter, J.C., Hutchison, C.A., and Smith, H.O. (2009). Enzymatic assembly of DNA molecules up to several hundred kilobases. Nat Methods 6, 343–345. 10.1038/nmeth.1318.

30. Ferrières, L., Hémery, G., Nham, T., Guérout, A.M., Mazel, D., Beloin, C., and Ghigo, J.M. (2010). Silent mischief: Bacteriophage Mu insertions contaminate products of Escherichia coli random mutagenesis performed using suicidal transposon delivery plasmids mobilized by broad-host-range RP4 conjugative machinery. Journal of Bacteriology 192, 6418–6427. 10.1128/JB.00621-10.

31. Donnenberg, M.S., and Kaper, J.B. (1991). Construction of an eae deletion mutant of enteropathogenic Escherichia coli by using a positive-selection suicide vector. Infect Immun 59, 4310–4317. 10.1128/iai.59.12.4310-4317.1991.

32. Choi, K.H., and Schweizer, H.P. (2006). mini-Tn7 insertion in bacteria with single attTn7 sites: Example Pseudomonas aeruginosa. Nature Protocols 1, 153–161. 10.1038/nprot.2006.24.

33. Duncan, S.H., Hold, G.L., Harmsen, H.J.M., Stewart, C.S., and Flint, H.J. (2002). Growth requirements and fermentation products of Fusobacterium prausnitzii, and a proposal to reclassify it as Faecalibacterium prausnitzii gen. nov., comb. nov. Int J Syst Evol Microbiol 52, 2141–2146. 10.1099/00207713-52-6-2141.

34. Shaw, P.D., Ping, G., Daly, S.L., Cha, C., Cronan, J.E., Rinehart, K.L., and Farrand, S.K. (1997). Detecting and characterizing *N*-acyl-homoserine lactone signal molecules by thin-layer chromatography. Proc. Natl. Acad. Sci. U.S.A. 94, 6036–6041. 10.1073/pnas.94.12.6036.

35. Zhu, J., Beaber, J.W., Moré, M.I., Fuqua, C., Eberhard, A., and Winans, S.C. (1998). Analogs of the Autoinducer 3-Oxooctanoyl-Homoserine Lactone Strongly Inhibit Activity of the TraR Protein of *Agrobacterium tumefaciens*. J Bacteriol 180, 5398–5405. 10.1128/JB.180.20.5398-5405.1998.

36. Bhagwat, A.A., Ying, Z.I., Karns, J., and Smith, A. (2013). Determining RNA quality for NextGen sequencing: some exceptions to the gold standard rule of 23S to 16S rRNA ratios. Microbiol Discov 1, 10. 10.7243/2052-6180-1-10.

37. Dobin, A., Davis, C.A., Schlesinger, F., Drenkow, J., Zaleski, C., Jha, S., Batut, P., Chaisson, M., and Gingeras, T.R. (2013). STAR: ultrafast universal RNA-seq aligner. Bioinformatics 29, 15–21. 10.1093/bioinformatics/bts635.

38. Andrews, S. (2010). FastQC: A Quality Control Tool for High Throughput Sequence Data [Online].

39. Ewels, P., Magnusson, M., Lundin, S., and Käller, M. (2016). MultiQC: Summarize analysis results for multiple tools and samples in a single report. Bioinformatics 32, 3047–3048. 10.1093/bioinformatics/btw354.

40. Wang, L., Wang, S., and Li, W. (2012). RSeQC: Quality control of RNA-seq experiments. Bioinformatics 28, 2184–2185. 10.1093/bioinformatics/bts356.

41. Love, M.I., Huber, W., and Anders, S. (2014). Moderated estimation of fold change and dispersion for RNA-seq data with DESeq2. Genome Biol 15, 550. 10.1186/s13059-014-0550-8.

42. Ignatiadis, N., Klaus, B., Zaugg, J.B., and Huber, W. (2016). Data-driven hypothesis weighting increases detection power in genome-scale multiple testing. Nat Methods 13, 577–580. 10.1038/nmeth.3885.

43. Zhu, A., Ibrahim, J.G., and Love, M.I. (2019). Heavy-tailed prior distributions for sequence count data: removing the noise and preserving large differences. Bioinformatics 35, 2084– 2092. 10.1093/bioinformatics/bty895.

44. Subramanian, A., Tamayo, P., Mootha, V.K., Mukherjee, S., Ebert, B.L., Gillette, M.A., Paulovich, A., Pomeroy, S.L., Golub, T.R., Lander, E.S., et al. (2005). Gene set enrichment analysis: A knowledge-based approach for interpreting genome-wide expression profiles. Proc. Natl. Acad. Sci. U.S.A. 102, 15545–15550. 10.1073/pnas.0506580102.

45. Mootha, V.K., Lindgren, C.M., Eriksson, K.-F., Subramanian, A., Sihag, S., Lehar, J., Puigserver, P., Carlsson, E., Ridderstråle, M., Laurila, E., et al. (2003). PGC-1α-responsive genes involved in oxidative phosphorylation are coordinately downregulated in human diabetes. Nat Genet 34, 267–273. 10.1038/ng1180.

46. Wu, T., Hu, E., Xu, S., Chen, M., Guo, P., Dai, Z., Feng, T., Zhou, L., Tang, W., Zhan, L., et al. (2021). clusterProfiler 4.0: A universal enrichment tool for interpreting omics data. The Innovation 2, 100141. 10.1016/j.xinn.2021.100141.

47. Taylor, S.C., Nadeau, K., Abbasi, M., Lachance, C., Nguyen, M., and Fenrich, J. (2019). The Ultimate qPCR Experiment: Producing Publication Quality, Reproducible Data the First Time. Trends in Biotechnology 37, 761–774. 10.1016/j.tibtech.2018.12.002.

48. Coulthrust, S.J., Clare, S., Evans, T.J., Foulds, I.J., Roberts, K.J., Welch, M., Dougan, G., and Salmond, G.P.C. (2007). Quorum sensing has an unexpected role in virulence in the model pathogen Citrobacter rodentium. EMBO Reports 8, 698–703. 10.1038/sj.embor.7400984.

49. Hense, B.A., and Schuster, M. (2015). Core Principles of Bacterial Autoinducer Systems. Microbiol Mol Biol Rev 79, 153–169. 10.1128/MMBR.00024-14.

50. Dong, T., and Schellhorn, H.E. (2010). Role of RpoS in Virulence of Pathogens. Infect Immun 78, 887–897. 10.1128/IAI.00882-09.

51. Spratt, M.R., and Lane, K. (2022). Navigating Environmental Transitions: the Role of Phenotypic Variation in Bacterial Responses. mBio, e02212–22. 10.1128/mbio.02212-22.

52. Dyszel, J.L., Smith, J.N., Lucas, D.E., Soares, J.A., Swearingen, M.C., Vross, M.A., Young, G.M., and Ahmer, B.M.M. (2010). *Salmonella enterica* Serovar Typhimurium Can Detect Acyl Homoserine Lactone Production by *Yersinia enterocolitica* in Mice. J Bacteriol 192, 29–37. 10.1128/JB.01139-09.

53. Landman, C., Grill, J.-P., Mallet, J.-M., Marteau, P., Humbert, L., Le Balc’h, E., Maubert, M.-A., Perez, K., Chaara, W., Brot, L., et al. (2018). Inter-kingdom effect on epithelial cells of the N-Acyl homoserine lactone 3-oxo-C12:2, a major quorum-sensing molecule from gut microbiota. PLoS ONE 13, e0202587. 10.1371/journal.pone.0202587.

54. Xue, J., Chi, L., Tu, P., Lai, Y., Liu, C.-W., Ru, H., and Lu, K. (2021). Detection of gut microbiota and pathogen produced N-acyl homoserine in host circulation and tissues. npj Biofilms Microbiomes 7, 53. 10.1038/s41522-021-00224-5.

55. Kim, M.K., Ingremeau, F., Zhao, A., Bassler, B.L., and Stone, H.A. (2016). Local and global consequences of flow on bacterial quorum sensing. Nat Microbiol 1, 15005. 10.1038/nmicrobiol.2015.5.

56. Siryaporn, A., Kim, M.K., Shen, Y., Stone, H.A., and Gitai, Z. (2015). Colonization, Competition, and Dispersal of Pathogens in Fluid Flow Networks. Current Biology 25, 1201–1207. 10.1016/j.cub.2015.02.074.

57. Arnoldini, M., Vizcarra, I.A., Peña-Miller, R., Stocker, N., Diard, M., Vogel, V., Beardmore, R.E., Hardt, W.-D., and Ackermann, M. (2014). Bistable Expression of Virulence Genes in Salmonella Leads to the Formation of an Antibiotic-Tolerant Subpopulation. PLoS Biol 12, e1001928. 10.1371/journal.pbio.1001928.

58. Gorelik, O., Levy, N., Shaulov, L., Yegodayev, K., Meijler, M.M., and Sal-Man, N. (2019). Vibrio cholerae autoinducer-1 enhances the virulence of enteropathogenic Escherichia coli. Sci Rep 9, 4122. 10.1038/s41598-019-40859-1.

59. Subramoni, S., and Venturi, V. (2009). LuxR-family ‘solos’: bachelor sensors/regulators of signalling molecules. Microbiology 155, 1377–1385. 10.1099/mic.0.026849-0.

60. Whiteley, M., Diggle, S.P., and Greenberg, E.P. (2017). Progress in and promise of bacterial quorum sensing research. Nature 551, 313–320. 10.1038/nature24624.

61. Paluch, E., Rewak-Soroczyńska, J., Jędrusik, I., Mazurkiewicz, E., and Jermakow, K. (2020). Prevention of biofilm formation by quorum quenching. Appl Microbiol Biotechnol 104, 1871–1881. 10.1007/s00253-020-10349-w.

62. Zhu, J., Miller, M.B., Vance, R.E., Dziejman, M., Bassler, B.L., and Mekalanos, J.J. (2002). Quorum-sensing regulators control virulence gene expression in Vibrio cholerae. Proceedings of the National Academy of Sciences of the United States of America 99, 3129– 3134. 10.1073/pnas.052694299.

63. Bridges, A.A., and Bassler, B.L. (2019). The intragenus and interspecies quorum-sensing autoinducers exert distinct control over Vibrio cholerae biofilm formation and dispersal. PLoS Biol 17, e3000429. 10.1371/journal.pbio.3000429.

64. Krzyżek, P. (2019). Challenges and Limitations of Anti-quorum Sensing Therapies. Front. Microbiol. 10, 2473. 10.3389/fmicb.2019.02473.

