## Supplementary Figures and Tables for "Quorum Sensing Modulates Bacterial Virulence and Colonization Dynamics During an Enteric Infection"

*Correspondence and requests for materials should be addressed to B.B.F.

### Supplemental Figures and Tables:

**Table S1:** Bacterial strains used in this study.

| Strain | Description | Source |
|---|---|---|
| DBS100 | *C. rodentium* ATCC 51459 | Schauer and Falkow[1] |
| DBS100 Δ*croI* | Acyl-homoserine-lactone synthase mutant | (Croxen and Koymans, unpublished) |
| DBS100 Δ*croR* | LuxR-type AHL receptor mutant | This study |
| DBS100 Δ*croI::croI* | Chromosomal complementation of Δ*croI* | This study |
| DBS100 Δ*croR::croR* | Chromosomal complementation of Δ*croR* | This study |
| MFD*pir* | *E. coli* conjugal donor for biparental matings (DAP auxotroph) | Ferrières et al.[2] |
| pTNS2 | *E. coli* with a Tn7 transposase expression plasmid ($Amp^R$) | Choi and Schweizer[3] |

**Table S2:** Plasmids used in this study.

| Plasmids | Description | Source |
|---|---|---|
| pRE112 | Suicide vector for allelic exchange ($Cam^R$) | Edwards et al.[4] |
| pUC18R6KT-mini-Tn7T | Mini-Tn7 base vector used for constructing chromosomally complemented strains ($Amp^R$) | Choi and Schweizer[3] |

**Table S3:** Oligonucleotide primers used for construction of mutant strains.

| Primer Name | Sequence* | Notes |
|---|---|---|
| ΔcroI1_FW | GTACGGTACCGAAGCTCTTGCGCCAT | KpnI |
| ΔcroI1_RV | GTACGCATGCTTCTCTTGTCAGGCGG | SphI |
| ΔcroI2_FW | ATGAACTCTGTTGTTGAGTTTCAACAA | |
| ΔcroI2_RV | AAACTCAACAACAGAGTTCATTG | |
| ΔcroR1_FW | aagcggtgtaagtgaactgcatgaattcccgggagagctcACATA<br>GGTGGGTTTAGCATACTGGAAGATGATGGG | SacI |
| ΔcroR1_RV | GGGGCTCAGTTATCCAGTAAGTACGTATCCTT<br>CACGTCAGTTACAGCCT | |
| ΔcroR2_FW | CTGACGTGAAGGATACGTACTTACTGGATAA<br>CTGAGCCCCGGATACCTGC | |
| ΔcroR2_RV | ggcccgatcccaagcttcttctagaggtaccgcatgcgatAAGGA<br>GGGGTGTTGCAGAGTGGATTATATGAATCT | KpnI |
| pRE112_FW | atcgcatgcggtacctctagaagaagcttg | Confirming fragment insertion in pRE112 |
| pRE112_RV | gagctctcccgggaattcatgcagttcac | Confirming fragment insertion in pRE112 |

*Restriction enzyme sites are underlined

**Table S4:** Oligonucleotide primers used for the construction of chromosomally complemented strains.

| Primer Name | Sequence* | Notes |
|---|---|---|
| Tn7-CroI_FW | gccttcgcgaggtaccTGTACCCGAGAGTACAAATGTAAGATAATCCCGTGCG | KpnI |
| Tn7-CroI_RV | ggctgcaggaattcctcgagTTACATACGGCGCAGTTGTTGAAGCATCTCACG | XhoI |
| Tn7-CroR_FW | gccttcgcgaggtaccCGGGATTTCTAAAGAAGAAGTTTGTACATTGATTAATGATATAAGCCT | KpnI |
| Tn7-CroR_RV | ggctgcaggaattcctcgagTCAGTTATCCAGTAATCTGAGCTCCATGCCGAGTTTTACCGCATG | XhoI |
| PTn7R | CACAGCATAACTGGACTGATTTC | Confirming chromosomal insertion of mini-Tn7 elements |
| PglmS-down_Citro | GCACGTTGAGGAAGTCATTGC | Confirming chromosomal insertion of mini-Tn7 elements |
| Tn7-check_FW | tagttgggaactgggagggg | Confirming fragment insertion in mini-Tn7 |
| Tn7-check_RV | tccgaagttcctattctctagaaagt | Confirming fragment insertion in mini-Tn7 |

* Restriction enzyme sites are underlined

**Table S5:** Olizgonucleotide primers used in this study for RT-qPCR.

| Primer Name | Sequence | Notes |
|---|---|---|
| dnaQ_FW | GTCAGGCCCGCAAATTAC | Endogenous control |
| dnaQ_RV | TCTCCACCAGATCGAGACG | Endogenous control |
| recA_FW | TGCCACTACCTGGCTGAAAG | Endogenous control |
| recA_RV | CGTGGAGTCCTGGTTGTTGA | Endogenous control |
| eae_FW | GGTTAATCTGCAGAGCGGTAA | Intimin (Type III secretion system) |
| eae_RV | GAACGGTAATAAGAAGTCCAGTGAA | Intimin (Type III secretion system) |
| espA_FW | AATCACCGGCGCTTAACTCA | Translocator protein (Type III secretion system) |
| espA_RV | TTCACGCACAAAGCGAACTG | Translocator protein (Type III secretion system) |
| espB_FW | ATGTTCGCAGATCGCAGGAT | Effector protein (Type III secretion system) |
| espB_RV | TTACCCTGCTAAACGAGCCG | Effector protein (Type III secretion system) |
| tir_FW | TTGCATCGACCCAATGGTCA | Intimin receptor (Type III secretion system) |
| tir_RV | AGTGCTTTGGATACCCTGCC | Intimin receptor (Type III secretion system) |
| rpoS_FW | GAAGACACCACGCAGGATGA | RNA polymerase sigma factor (Stress Response) |
| rpoS_RV | CCGCTTCATAACCCAGCAGA | RNA polymerase sigma factor (Stress Response) |
| katE_FW | CATGACACCCGTGAATCCCA | Catalase HPII (Stress Response) |
| katE_RV | CCGTATCCGGCGATTTCAGA | Catalase HPII (Stress Response) |
| croI_FW | GCAGCCAGTCACCAGCTTTG | AHL Synthase (Quorum Sensing) |
| croI_RV | TGCGAACTGGAAGGTGGAGG | AHL Synthase (Quorum Sensing) |

**Table S6: MRM parameters optimized for the identification and quantification of AHLs.** Multiple-reaction monitoring (MRM) parameters used for AHL detection and quantification. MF, Molecular Formula; MW, Molecular Weight; Q1, Precursor Ion; Q3, Product Ion; Rt, Retention Time; Frag, Fragmentor; CE, Collision Energy.

| Compound | MF | MW | Q1/Q3 quantification | Q1/Q3 confirmation | Rt (min) | Dwell | Frag (V) |
|---|---|---|---|---|---|---|---|
| C4-HSL | $C_8H_{13}NO_3$ | 171.0895 | 172.0/102 | 172/43 | 8.4 | 80 | 80 |
| C6-HSL | $C_{10}H_{17}NO_3$ | 199.1208 | 200.1/102 | 200.1/71 | 12.0 | 80 | 80 |
| 3-oxo-C6-HSL | $C_{10}H_{15}NO_4$ | 213.1001 | 214.1/113 | 214.1/102 | 9.1 | 60 | 70 |
| 3-hydroxy-C6-HSL | $C_{10}H_{17}NO_4$ | 215.1158 | 216.1/102 | 216.1/69 | 8.0 | 60 | 80 |

| Compound | CE (V) | Cell Accelerator (V) |
|---|---|---|
| C4-HSL | 5 | 7 |
| C6-HSL | 6 | 7 |
| 3-oxo-C6-HSL | 5 | 7 |
| 3-hydroxy-C6-HSL | 5 | 7 |

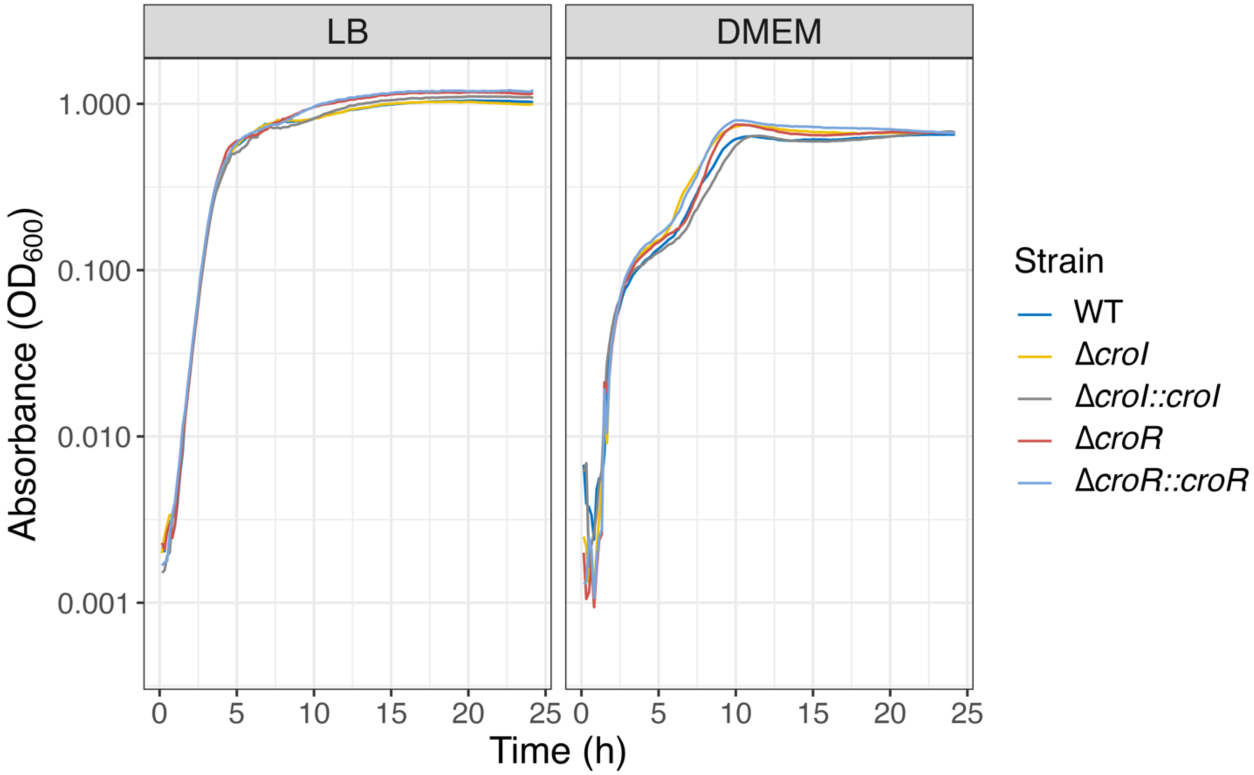


**Figure S1: *In vitro* growth of *C. rodentium* of WT DBS100, Δ*croI*, Δ*croR*, and complemented strains.**

*C. rodentium* strains grown over the course of 24 hours in either LB or DMEM media. Cultures were grown at 37 °C with continuous shaking. Curves represent the average of three biological replicates measured in three technical replicates each. LB, lysogeny broth. DMEM, Dulbecco's Modified Eagle Medium.

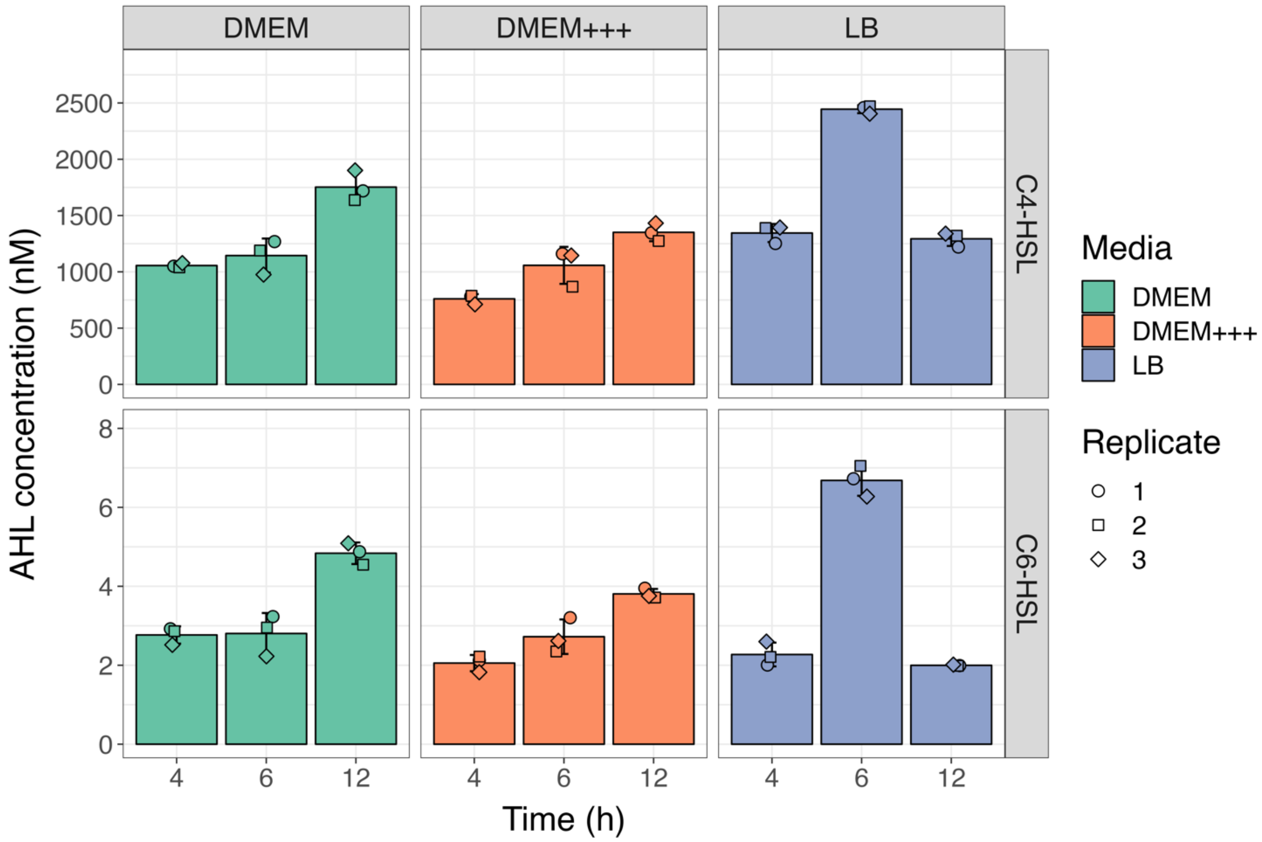


**Figure S2: Accumulation of C4-HSL and C6-HSL produced by *C. rodentium* during growth in different culture media.**
C4-HSL and C6-HSL produced by *C. rodentium* over time in liquid culture, measured in cell-free supernatants collected and partially purified using ethyl acetate before analysis via LC-MS/MS. Trace amounts of 3-hydroxy-C6-HSL were also detected in all media (data not shown). Data represent the mean of 3 biological replicates ± SD. DMEM, Dulbecco's Modified Eagle Medium. DMEM+++, Dulbecco's Modified Eagle Medium supplemented with 10 % heat-inactivated fetal bovine serum, 1 % Non-Essential Amino Acids, and 1 % L-glutamine. LB, lysogeny broth.

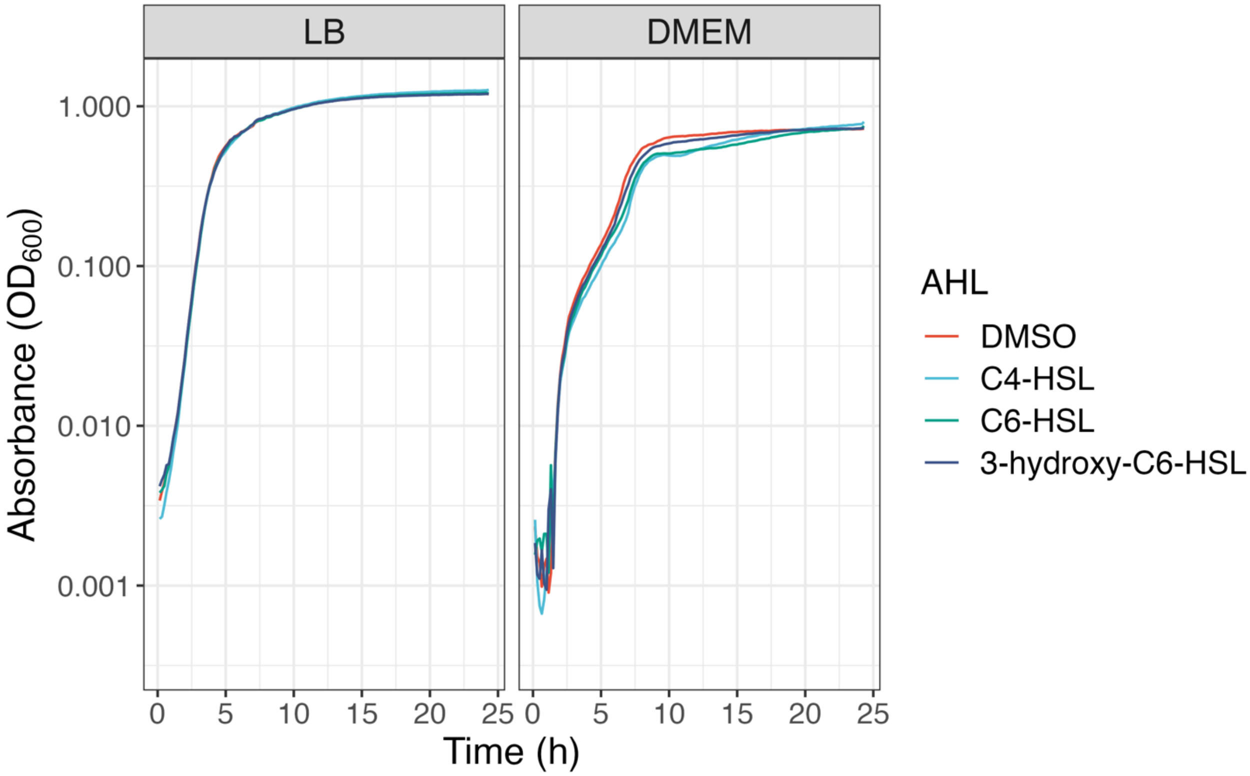


**Figure S3: Growth dynamics of WT *C. rodentium* supplemented with different AHLs.**
WT *C. rodentium* was grown in LB or DMEM media for 24 hours at 37 °C with continuous shaking. Cultures were supplemented with either 10 µM of C4-HSL, C6-HSL, 3-hydorxy-C6-HSL or DMSO as vehicle control, showing no impact of AHLs on growth of *C. rodentium*. Data represent the average of three independent biological replicates with three technical replicates each.

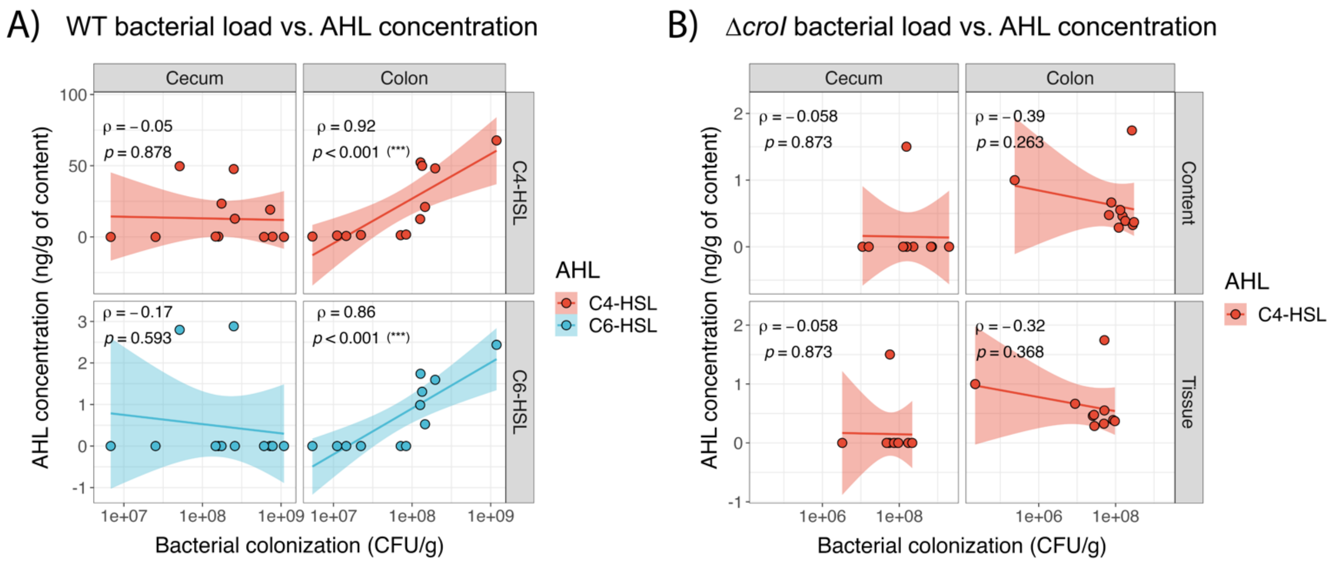


**Figure S4: *In vivo* AHL concentration measured in the colon correlates with *C. rodentium* burden.**

**A)** Spearman correlations between the detected C4-HSL and C6-HSL concentration and the CFU burden of WT *C. rodentium* in cecal and colonic lumenal content at day 7 post-infection. **B)** Spearman correlations of detected C4-HSL concentrations and the CFU burden of Δ*croI C. rodentium* in lumenal and tissue-associated subpopulations at the cecum and colon on day 7 post-infection. Rho ($\rho$) represents the Spearman correlation values; ($p$) represents the $p$-value. Line represents the linear regression; shaded area represents the 95 % confidence interval.

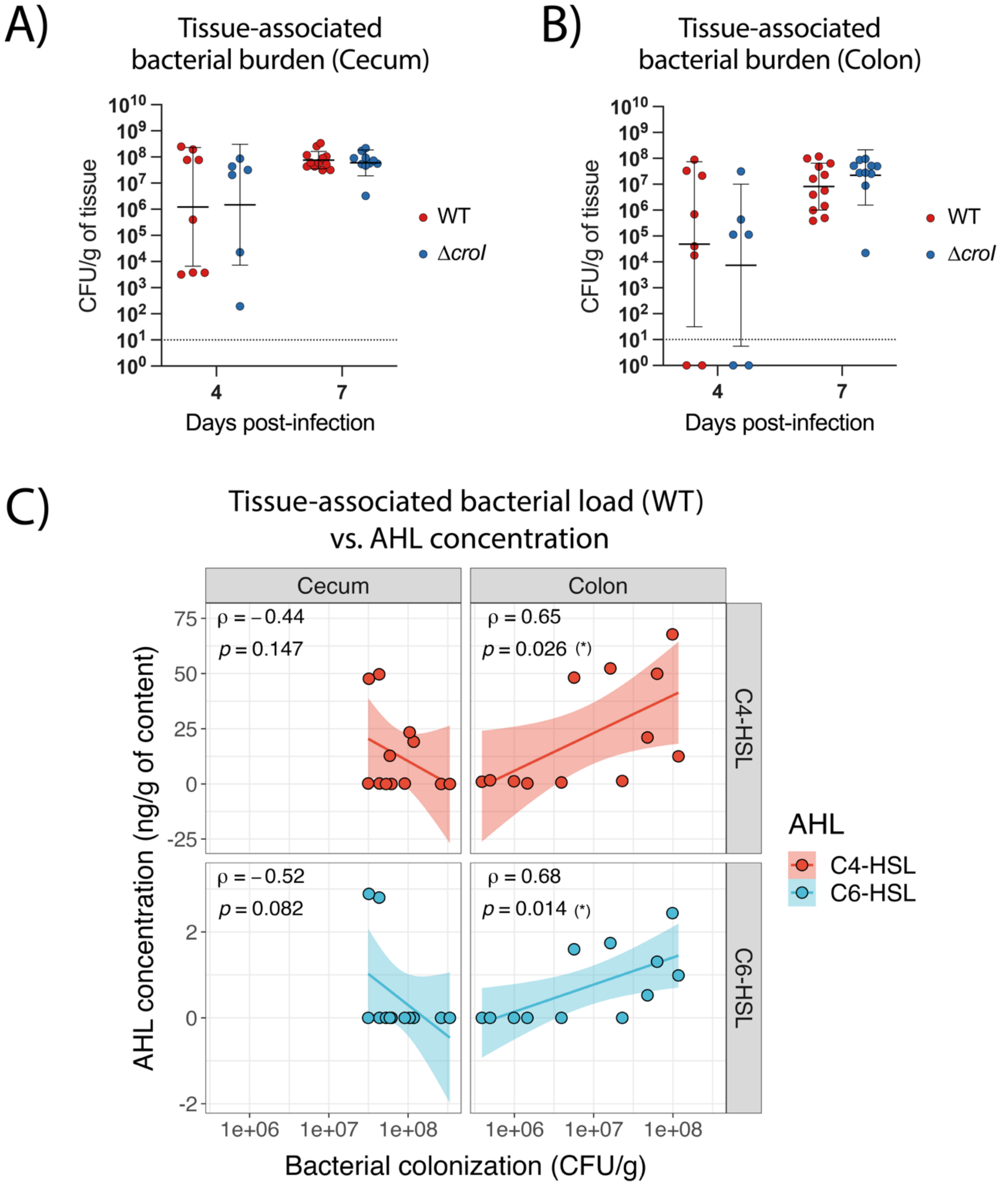


**Figure S5: Mucosal-associated CFU burden of *C. rodentium* during infection.**
**A-B)** Tissue-associated burden of WT *C. rodentium* in the cecum and colon at day 4 and 7 post-infection (p.i.). Lines represent geometric mean ± geometric standard deviation, with an N = 6-12 mice per group. The limit of detection is displayed as a dotted line. **C)** Spearman correlations between the detected C4-HSL and C6-HSL concentration and the tissue-associated burden of WT *C. rodentium* in the cecum and colon at day 7 post-infection. Rho ($\rho$) represents the Spearman correlation values; ($p$) represents the $p$-value. Line represents the linear regression; shaded area represents the 95 % confidence interval.

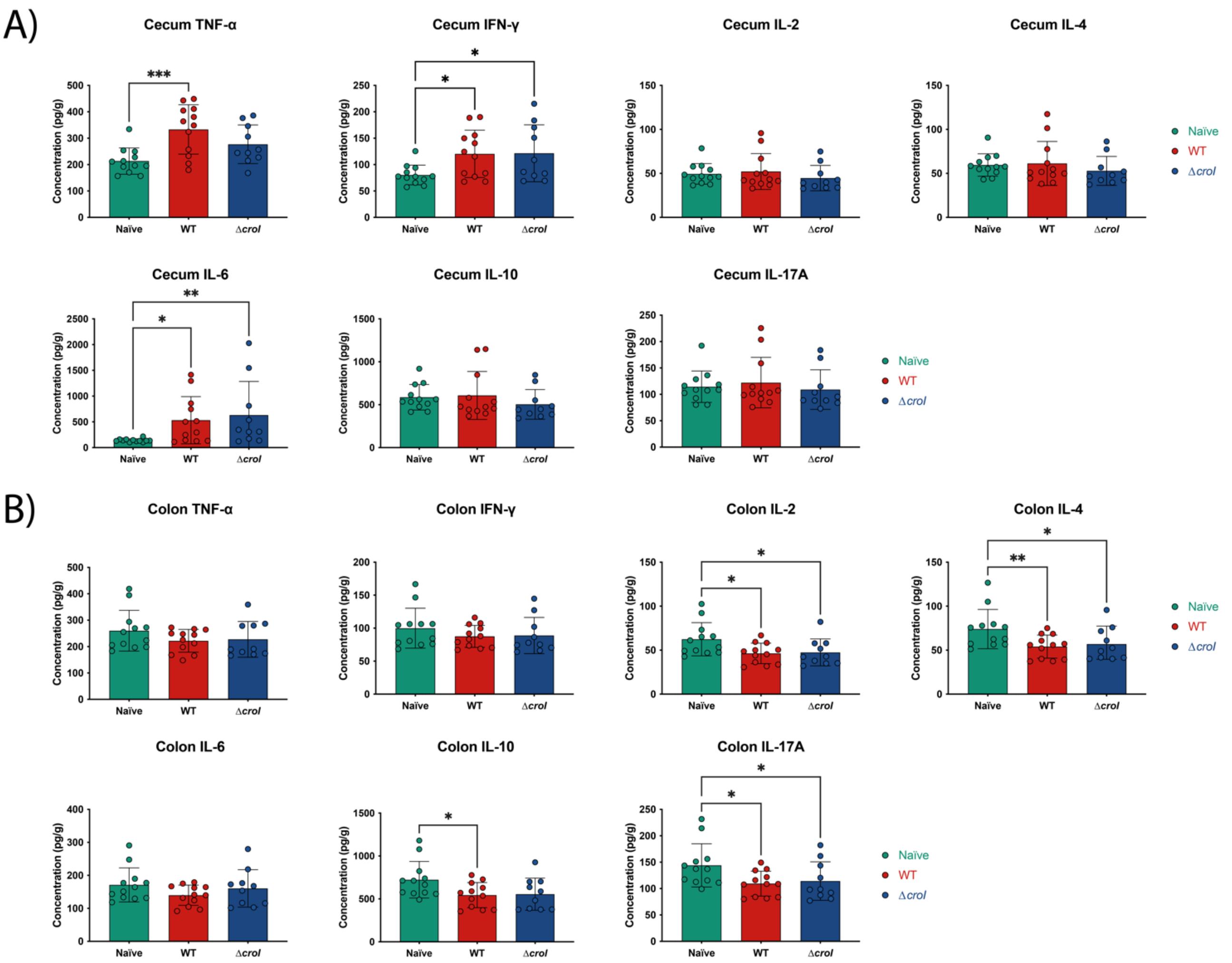


**Figure S6: Measurement of inflammatory cytokine markers in cecal and colonic tissues of uninfected and WT- and Δ*croI*-infected mice.**

Tissues were collected at day 7 post infection and were measured using Cytokine Bead Array (CBA). Data represent the mean ± SD with an N = 10-12. Statistical analysis was performed by using a Kruskal-Wallis test. TNF-$\alpha$, Tumour Necrosis Factor alpha; IFN-$\gamma$, Interferon gamma; IL-2, Interleukin-2; IL-4, Interleukin-4; IL-6, Interleukin-6; IL-10, Interleukin-10; IL-17, Interleukin-17.

### References:


1. Schauer, D.B., and Falkow, S. (1993). The eae gene of Citrobacter freundii biotype 4280 is necessary for colonization in transmissible murine colonic hyperplasia. Infection and Immunity *61*, 4654–4661. 10.1128/iai.61.11.4654-4661.1993.

2. Ferrières, L., Hémery, G., Nham, T., Guérout, A.M., Mazel, D., Beloin, C., and Ghigo, J.M. (2010). Silent mischief: Bacteriophage Mu insertions contaminate products of Escherichia coli random mutagenesis performed using suicidal transposon delivery plasmids mobilized by broad-host-range RP4 conjugative machinery. Journal of Bacteriology *192*, 6418–6427. 10.1128/JB.00621-10.

3. Choi, K.H., and Schweizer, H.P. (2006). mini-Tn7 insertion in bacteria with single attTn7 sites: Example Pseudomonas aeruginosa. Nature Protocols *1*, 153–161. 10.1038/nprot.2006.24.

4. Edwards, R.A., Keller, L.H., and Schifferli, D.M. (1998). Improved allelic exchange vectors and their use to analyze 987P fimbria gene expression. Gene *207*, 149–157. 10.1016/S0378-1119(97)00619-7.
